# Concerted evolution reveals co-adapted amino acid substitutions in Na^+^K^+^-ATPase of frogs that prey on toxic toads

**DOI:** 10.1101/2020.08.04.234435

**Authors:** Shabnam Mohammadi, Lu Yang, Arbel Harpak, Santiago Herrera-Álvarez, María del Pilar Rodríguez-Ordoñez, Julie Peng, Karen Zhang, Jay F. Storz, Susanne Dobler, Andrew J. Crawford, Peter Andolfatto

**Affiliations:** School of Biological Sciences, University of Nebraska, Lincoln, NE, USA; Department of Ecology and Evolutionary Biology, Princeton University, Princeton, NJ, USA; Department of Biological Sciences, Columbia University, New York, NY, USA; Department of Biological Sciences, Universidad de los Andes, Bogotá, 111711, Colombia; Lewis-Sigler Institute, Princeton University, Princeton, NJ, USA; Molecular Evolutionary Biology, Zoological Institute, Universität Hamburg, Hamburg, Germany

## Abstract

Gene duplication is an important source of evolutionary innovation, but the adaptive division-of-labor between duplicates can be opposed by ongoing gene conversion between them. Here we document a tandem duplication of Na^+^,K^+^-ATPase subunit α1 (ATP1A1) shared by frogs in the genus *Leptodactylus*, a group of species that feeds on toxic toads. One ATP1A1 paralog evolved resistance to toad toxins while the other paralog retained ancestral susceptibility. We show that the two *Leptodactylus* paralogs are distinguished by 12 amino acid substitutions that were maintained by strong selection that counteracted the homogenizing effect of gene conversion. Protein-engineering experiments show that two major-effect substitutions confer toxin resistance, whereas the 10 additional substitutions mitigate deleterious pleiotropic effects on enzyme function. Our results highlight how trans-specific, neofunctionalized gene duplicates can provide unique insights into interactions between adaptive substitutions and the genetic backgrounds on which they arise.

**One Sentence Summary:** Selection counteracts gene conversion to maintain an adaptive division-of-labor between tandemly duplicated genes.

## Main Text

The repeated evolution of toxin resistance in animals is one of the clearest examples of natural selection at the molecular level and represents a useful paradigm to examine constraints on the evolution of novel protein functions (*1*). Neotropical Grass Frogs of the genus *Leptodactylus* (Leptodactylidae) are widely distributed throughout lowland South America and are known to feed on chemically-defended toads – a predatory tendency that is rare among frogs (*2*–*6*). A major component of the chemical defense secretions of toads is a class of cardiac glycosides (CGs) called “bufadienolides” (*7*) that inhibit the α-subunit of Na^+^,K^+^-ATPases (ATP1A). Na^+^,K^+^-ATPases are transmembrane proteins that are vital to numerous physiological processes in animals including neural signal transduction, muscle contraction, and cell homeostasis (*8, 9*). CGs bind to the extracellular surface of ATP1A and block the flux of ions (*10*), making them potent poisons to most animals. However, some vertebrates have independently evolved the ability to prey on chemically-defended toads, partly via amino acid substitutions to the CG-binding domain of ATP1A1 that confer resistance to CGs (*11*–*14*).

Most vertebrates share several paralogous copies of ATP1A that have different tissue-specific expression profiles (*15*). For example, ATP1A1 is the most ubiquitously expressed paralog and ATP1A3 has enriched expression in nervous tissue and heart muscle (*16, 17*; Fig. S1). Previous studies on the molecular convergence of CG-resistance in reptiles have focused primarily on the αM1–2 extracellular loop of ATP1A3 (*12*–*14, 18*), whereas studies of birds, mammals, and amphibians have focused on the same region of ATP1A1 (*11, 18*). A survey of ATP1A1 αM1–2 in toads and frogs (*11*) revealed a possible duplication of this gene in the toad-eating frog, *Leptodactylus latrans* (reported as *L*. *ocellatus*), where the resistant (R) paralog includes substitutions known to confer resistance to CGs while the sensitive (S) paralog appears to have retained the ancestral susceptibility to CGs. Neofunctionalization of ATP1A paralogs has contributed to the evolution of CG-resistance in numerous insect lineages (*19*–*22*) but appear to be rare among CG-resistant vertebrates. Further, the fate of duplicated genes and the probability that they will neofunctionalize is predicted to depend on the strength of selection for functional differentiation relative to the rate of non-allelic gene conversion (NAGC), a form of nonreciprocal genetic exchange that homogenizes sequence variation between duplicated genes, thereby impeding divergence (*23*–*25*). The ATP1A1 duplication in *Leptodactylus* provides an ideal opportunity to explore this process because the functional differentiation between R and S paralogs has clear adaptive significance with regard to CG-resistance.

We surveyed the full-length coding sequences of all ATP1A paralogs in *Leptodactylus* and other anurans using RNA-seq-based gene discovery (*19*; Tables S1). Our results confirm that ATP1A1 is duplicated in *L*. *latrans* (*11*) and indicate that the duplication of ATP1A1 most likely occurred in the common ancestor of all surveyed *Leptodactylus* species (Fig. 1A, Table S2). Two other ancient paralogs common to vertebrates, ATP1A2 and ATP1A3, appear to be present as single-copy genes and lack any known CG-resistant substitutions in *Leptodactylus* (Fig. S2). The αM1– 2 transmembrane domains of the ATP1A1 paralogs in *L*. *latrans* are distinguished by four amino acid substitutions (*11*; Fig 1C, Fig. S3). Two of these substitutions, Q111R and N122D, were first identified in rat ATP1A1 and have been shown to interact synergistically to confer CG-resistance to sheep ATP1A1 protein *in vitro* (*26, 27*). Comparison of ATP1A1 sequences in five distantly related *Leptodactylus* species (*28*) reveals that they each harbor a putatively resistant paralog (R) that includes the Q111R and N122D substitutions and a putatively sensitive ATP1A1 paralog (S) that lacks these substitutions. In addition to Q111R and N122D, there are 10 other amino acid substitutions distinguishing the R and S paralogs in most of the five sampled species (Fig. 1C). Hereafter, we refer to these twelve substitutions as “R/S distinguishing”.

**Fig. 1.**
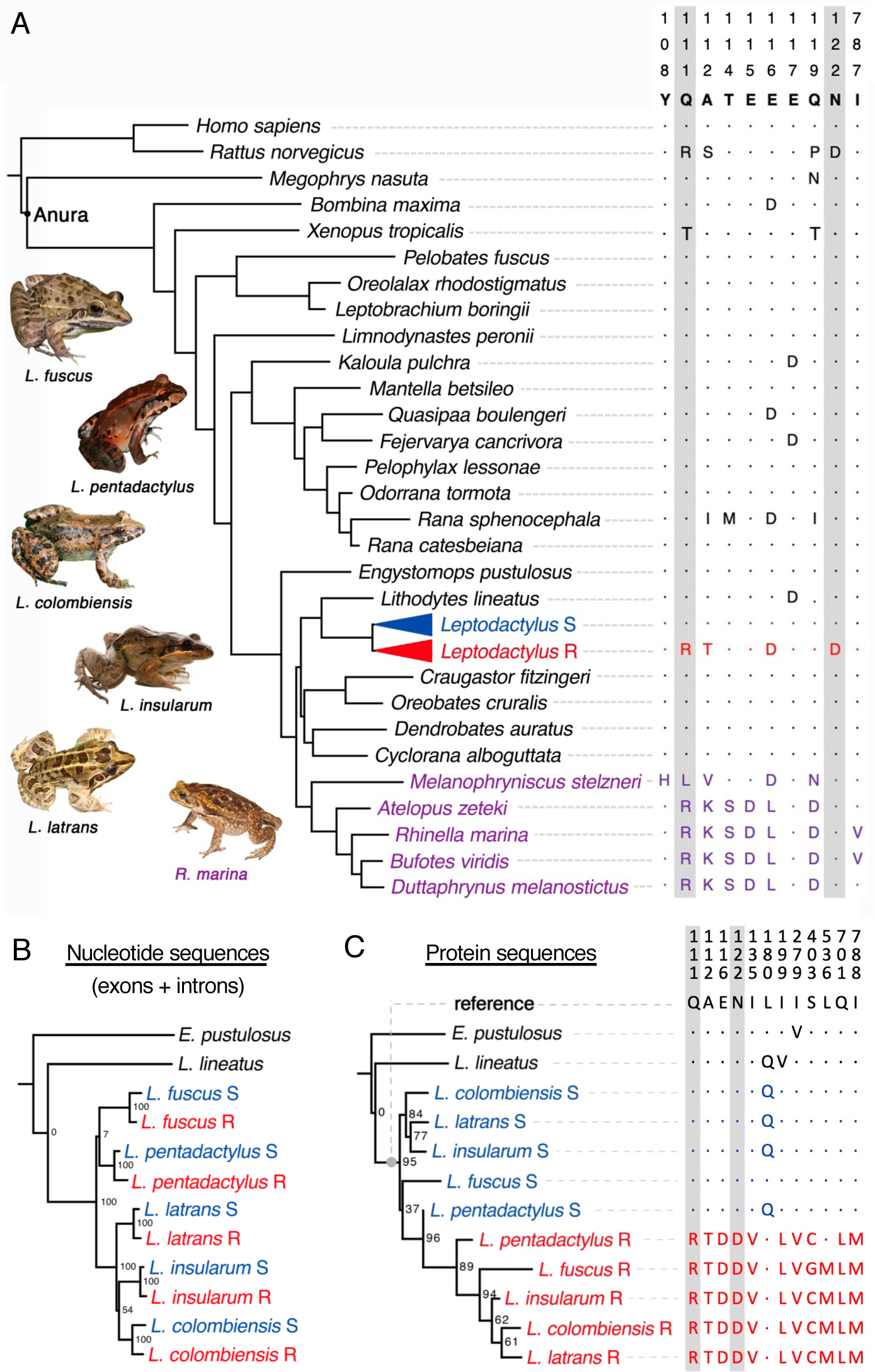
Molecular evolution of ATP1A1 in anurans. (A) Maximum likelihood phylogeny of anuran species and mammalian outgroups derived from (*38*). Species names in purple correspond to chemically defended toads, and blue and red colors correspond to the S and R ATP1A1 paralogs in *Leptodactylus* species, respectively. Only variable sites with documented roles in CG-binding or sensitivity are shown (reviewed in (*22*)). The numbering of sites is based on sheep ATP1A1 (*Ovis aries*, Genbank: NC019458.2) and appears at the top of the table (e.g. the first position shown is 108). Dots indicate identity with the reference sequence and letters represent amino acid substitutions relative to the reference. The images on the left depict the five surveyed *Leptodactylus* species and a representative toad species (*Rhinella marina*) as potential prey. Maximum likelihood phylogeny estimates based on nucleotide sequences (B) and amino acid sequences (C) yield distinct topologies. Bootstrap support values are indicated at internal nodes. To the right is the pattern of amino acid variation at 12 positions that distinguish the S and R paralogs.

To infer when duplication occurred relative to speciation events, we estimated phylogenies from an alignment of ATP1A1 coding sequence. Phylogenies estimated from nucleotide and inferred amino-acid sequences support strikingly different topologies (Fig 1B and C). Without further information, the genealogy based on nucleotide sequences (primarily based on variation at synonymous sites) would imply independent duplications in each of the *Leptodactylus* species, followed by parallel substitutions at the same 12 R/S distinguishing amino acid positions (Fig. 1B). Instead, it seems much more likely that this genealogical pattern reflects a single ancestral duplication — as indicated by genealogy based on amino acid sequences (Fig. 1C, Table S4) — coupled with on-going NAGC between the R and S paralogs of each species. Frequent NAGC produces a pattern of “concerted evolution” whereby tandemly linked paralogs from the same species are more similar to one another than they are to their orthologous counterparts in other species (*29;* Fig. 2A,B). By generating a *de novo* genome assembly of *L*. *fuscus* based on single-molecule sequencing, we established that S and R copies are indeed arranged in tandem and in the same orientation, and are therefore likely to be subject to NAGC (Table S3, Fig. S4). We thus propose that the persistence of the 12 amino acid differences between the two paralogs is due to selection counteracting the homogenizing effects of NAGC (*23, 30*; Fig. 2B), thereby maintaining an adaptive division-of-labor between the R and S copies.

**Fig. 2.**
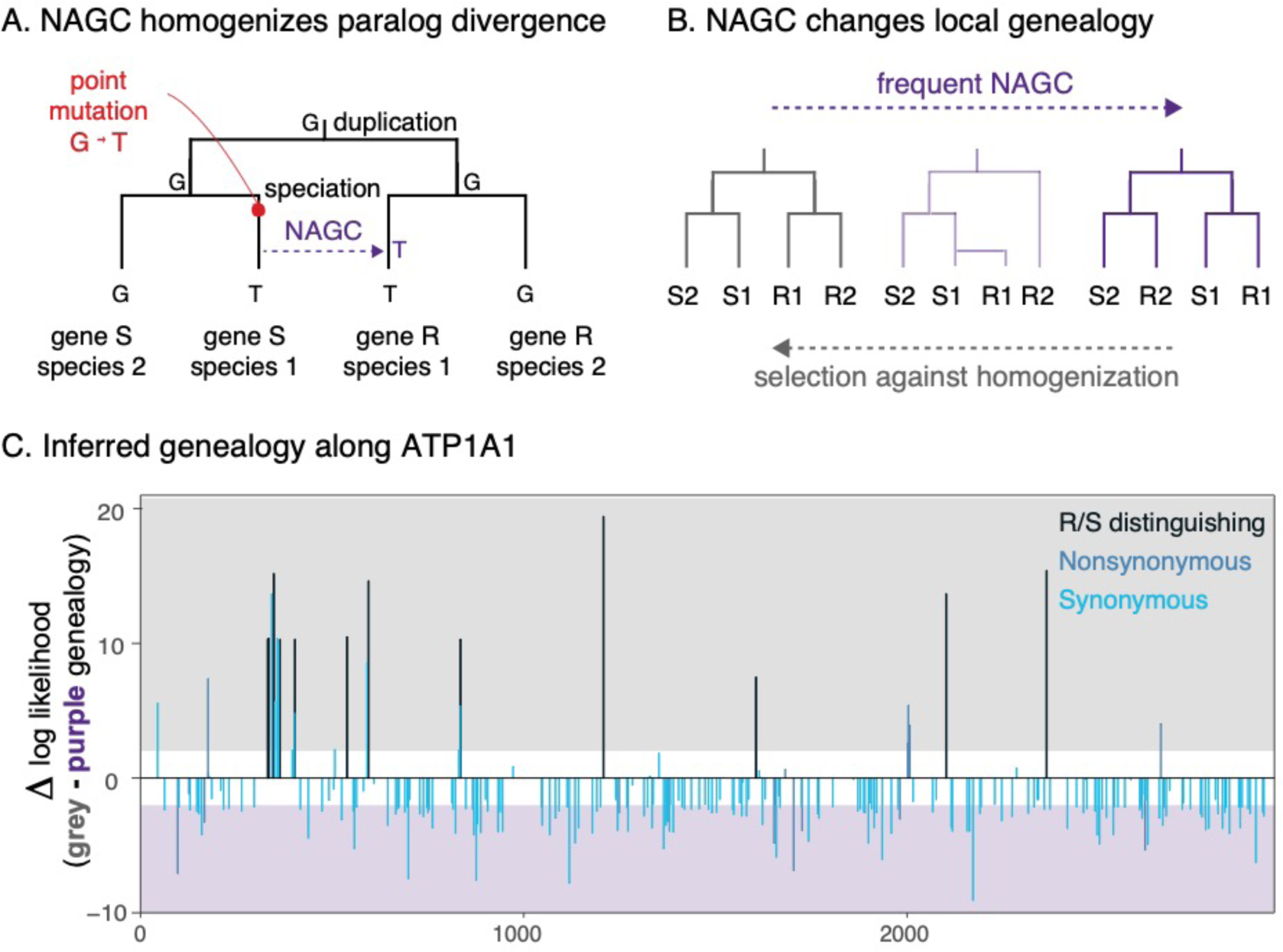
**(A)** Non-allelic gene conversion (NAGC) homogenizes sequence variation between paralogous genes, and therefore changes the genealogical signal (adapted from (*39*)). **(B)** NAGC can result in a genealogy in which paralogous genes in the same species share a more recent common ancestor with one another than with their orthologous counterparts in other species (“concerted evolution”). The homogenizing effects of NAGC can be counteracted by selection that favors the differentiation of paralogous genes. **(C)** Site-wise difference in the log_10_-likelihood of two alternative tree topologies—generalizing the grey and purple extremes of panel B to five *Leptodactylus* species. Shaded regions show a log-likelihood difference greater than 2 in support of the corresponding model. Only parsimony-informative variants in the ATP1A1 coding sequence are shown. Black bars correspond to the 12 R/S distinguishing nonsynonymous substitutions (shown in red or blue in Fig 1C; Table S4).

The opposing forces of NAGC and selection are predicted to leave a characteristic genealogical signature at neutral sites closely linked to the targets of selection (*30*; Fig. 2B). We therefore tested the relationship between the genealogical signature and distance from nonsynonymous variants putatively under selection. To this end, for all informative sites, we evaluated the level of support for an ancient duplication of ATP1A1 in the common ancestor of all *Leptodactylus* species (with no concerted evolution) relative to support for an alternative in which ATP1A1 paralogs within species are always more closely related to one another than they are to paralogs in other species (as expected under concerted evolution). This analysis reveals that synonymous (presumed to be neutral) variants congruent with an ancient duplication of R and S have a median distance of 4 bp from nonsynonymous variants exhibiting the same pattern (Fig. 2C). In contrast, randomly sampled synonymous sites supporting the alternative genealogy (i.e. concerted evolution) have a median distance of 88 bp from those nonsynonymous variants (bootstrap p<10^−5^). This pattern at synonymous sites is consistent with a scenario in which purifying selection maintains functionally important sequence differences between neofunctionalized gene duplicates in the face of NAGC.

We next quantified the strength of purifying selection required to maintain the amino acid differentiation between R and S duplicates in the face of NAGC. We first considered population genetics theory for the evolution of a single site in tandem duplicates (*31*) (Supplementary Materials). This analytic model predicts that if the rate of NAGC is an order of magnitude higher than the rate of point mutation, then the maintenance of alternative amino acid states is only likely under sufficiently strong purifying selection — namely, when the selection coefficient scaled by population size, 2*Ns*, is larger than one (Fig. 3A). We next developed an inference method based on simulations of ATP1A1 evolution to estimate the combination of parameters that best explain divergence patterns throughout the gene, including levels of paralog divergence observed as a function of distance from the 12 R/S distinguishing substitutions (Supplementary Materials). We estimate the rate of NAGC to be an order of magnitude higher than the point mutation rate (posterior mode 9 with 80% credible interval 4-54 times the point mutation rate), and 2*Ns* substantially larger than one (posterior mode 9; 80% credible interval 5-18; Fig. 3B). These estimates fall within the plausible range predicted by the theoretical single-site model (Fig. 3A). These results indicate that the observed pattern of divergence between R and S paralogs reflects a history of strong purifying selection that maintains fixed differences between them despite high rates of NAGC.

**Fig. 3.**
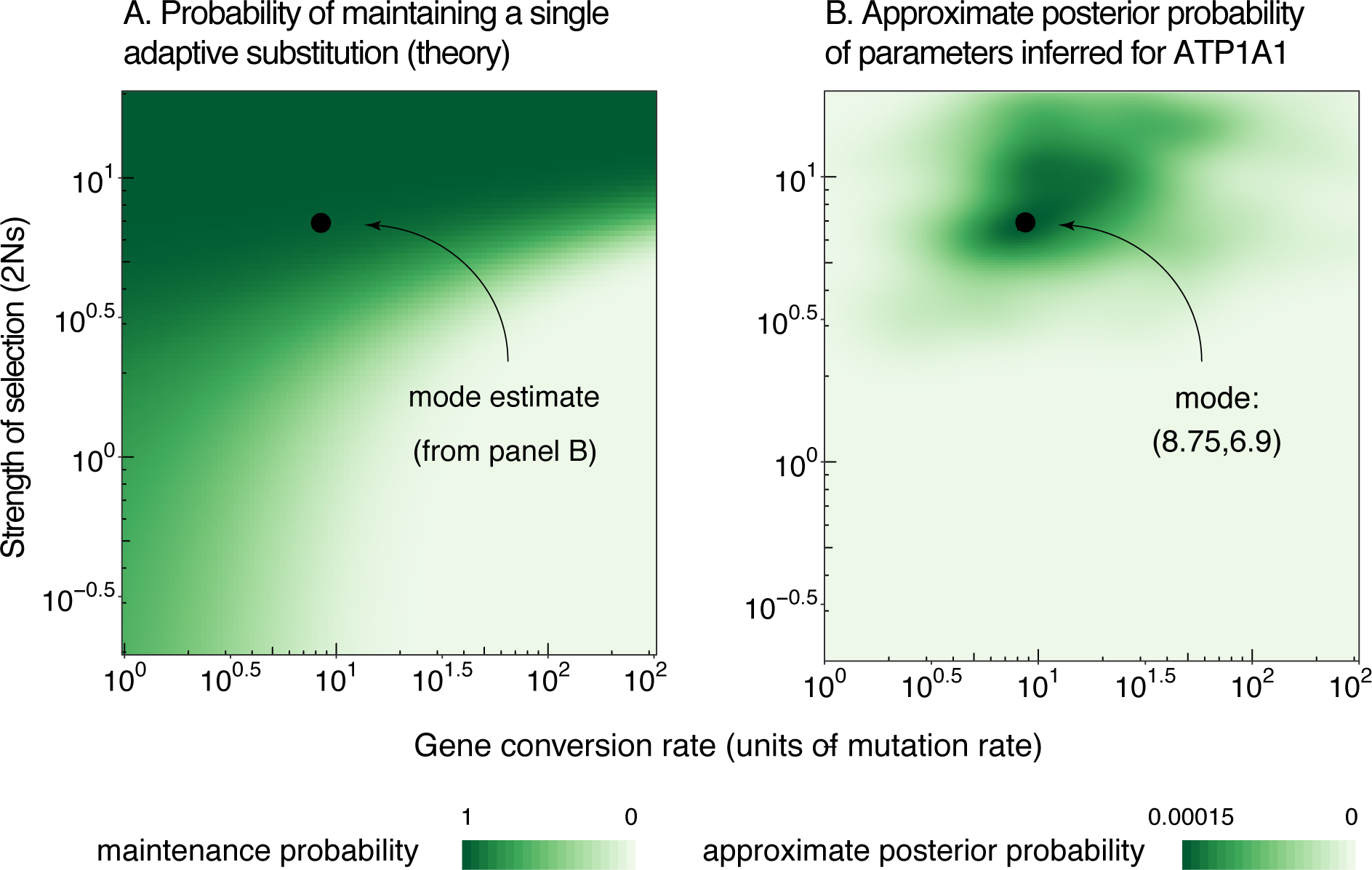
(A) Theoretical probability of maintaining distinct alleles at a single site in the face of non-allelic gene conversion (NAGC). We used a theoretical model to compute the probability of maintaining alternative amino acid states at the same site in a pair of paralogous genes, given an NAGC rate and strength of selection against allele homogenization at the site. The black dot shows the approximate mode estimate from panel B, which falls in the range in which maintenance is likely according to this theoretical model. **(B) Estimates of evolutionary parameters**. Approximate posterior probabilities were inferred based on simulations of the evolution of ATP1A1 genes in *Leptodactylus*. The x-axis shows the NAGC rate across the gene, and the y-axis shows the population selection coefficient for the 12 substitutions that distinguish the R and S paralogs across species.

The inference that selection maintains the co-occurrence of the 12 R/S distinguishing substitutions implies they are functionally important and collectively contribute to organismal fitness. The effects of Q111R and N122D on CG-insensitivity have previously been demonstrated by *in vitro* enzyme inhibition assays (*9*). Additionally, while not related directly to CG-resistance, the potential importance of substitutions at sites 112 and 116 has been suggested by molecular evolution analysis and structural studies, respectively (*11, 32*). However, the remaining eight R/S distinguishing substitutions are located in structural domains that have not been implicated in CG-resistance. Since our analysis suggests that amino acid divergence between R and S paralogs is maintained by selection, we performed protein-engineering experiments to elucidate the functional significance of the 12 R/S distinguishing substitutions. We synthesized and recombinantly expressed eight mutant Na^+^,K^+^-ATPase proteins, each harboring different combinations of R-specific replacements on both S- and R-type genetic backgrounds of a representative species, *L*. *latrans* (Fig. 4A, Table S6), and we then quantified the level of CG-resistance of each genotype using enzyme-inhibition assays (Table S7, Fig. S7, (*33*)). Individually, Q111R and N122D significantly increased CG-resistance by 21-fold and 14-fold, respectively (ANOVA p=2.7e-13 and p=2.3e-6; Fig. 4B, Table S8). When combined, Q111R and N122D produce a greater than 100-fold increase in CG resistance relative to the S paralog (Tukey’s HSD test, adjusted p<4e-5, Fig. 4B, Tables S6-S8). In contrast, the remaining 10 substitutions had no detectable net effect on CG-resistance when jointly added to the S background (p=0.22, Fig. 4B).

**Fig. 4.**
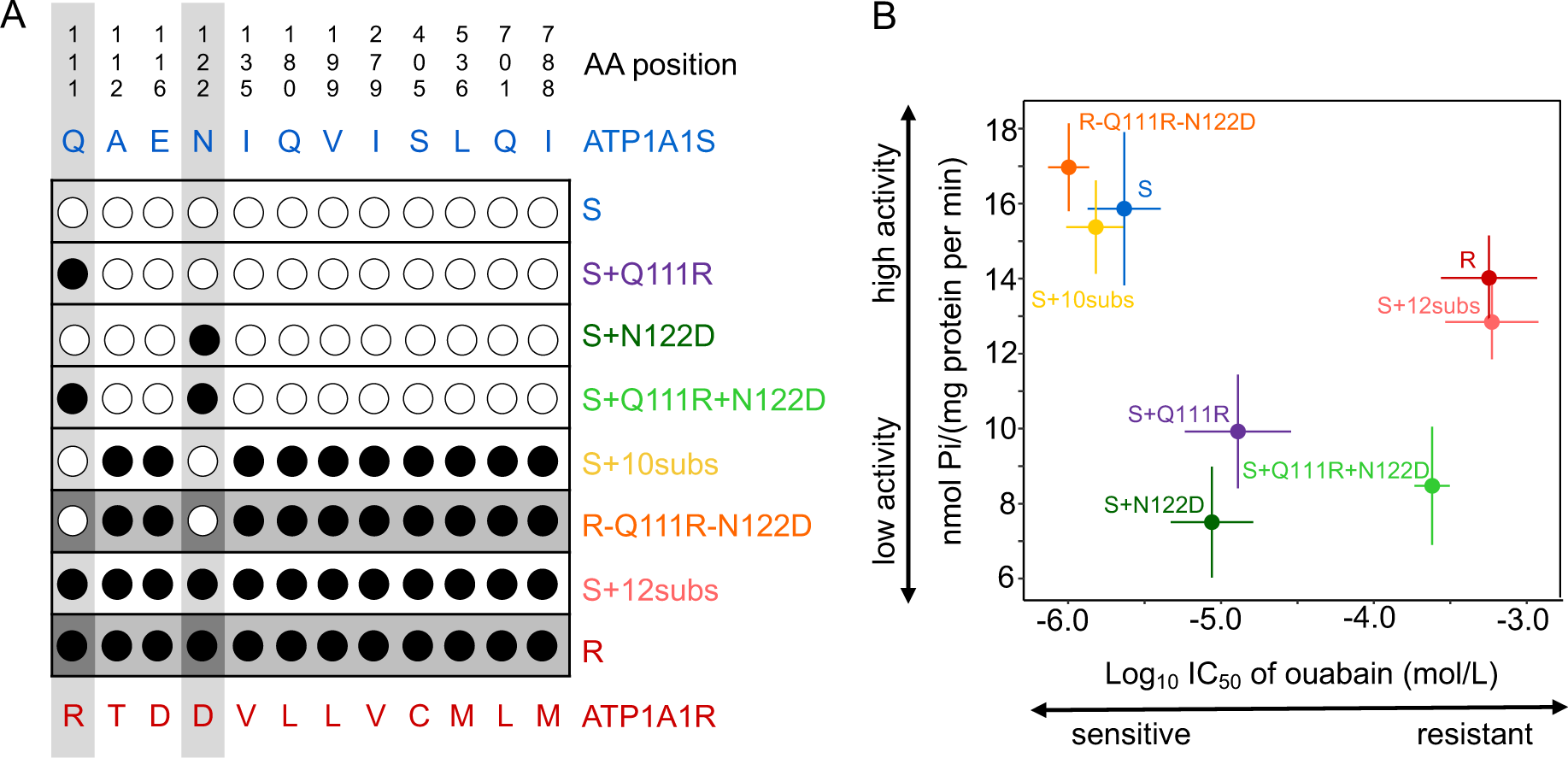
Functional analysis of substitutions specific to the R-type ATP1A1 paralog. (A) ATP1A1 gene constructs with various combinations of the 12 substitutions that distinguish the S and R paralogs. Black circles indicate an amino acid matching the R paralog whereas a white circle indicates a match with the S paralog. Dark grey shading denotes the R background and white denotes the S background. Light grey columns highlight two substitutions (Q111R and N122D) that are known to confer CG-resistance. (B) Functional properties of engineered Na^+^,K^+^-ATPases. The mean ± SEM log_10_IC_50_ (i.e., a measure of CG resistance) is plotted on the x-axis and the mean ± SEM ATP hydrolysis rate (i.e., a measure of protein activity) for the same proteins is plotted on the y-axis. Each estimate is based on six biological replicates.

Given the absence of detectable effects of R/S distinguishing substitutions other than Q111R and N122D on CG-resistance, we tested whether these substitutions had effects on other aspects of ATP1A1 function. Since ATP hydrolysis and ion co-transport are strongly coupled functions of Na^+^,K^+^-ATPase (*34*), we used estimates of the rate of ATP hydrolysis in the absence of ouabain as a proxy for overall protein activity. Based on this assay, we found that CG-resistance substitutions Q111R and N122D significantly impair activity, individually reducing ATPase activity by an average of 40% (*p*=0.024 and *p*=7.7e-4 respectively; Fig. 4B; Table S8). We also detected a significant interaction between Q111R and N122D that renders their joint effects somewhat less severe than predicted by the sum of their individual effects (*i*.*e*., a 30% reduction rather than the expected 78% reduction, p=0.022). Critically, adding the remaining 10 R-specific substitutions on the S background containing Q111R and N122D restores normal ATPase activity close to wild-type levels. An analysis of variance reveals a highly significant effect of these 10 R-specific substitutions (p=1.2 × 10^−4^, Fig. 4B, Table S8). Our results thus indicate that these 10 R/S distinguishing substitutions play a vital role in compensating for the negative pleiotropic effects of the resistance-conferring substitutions, Q111R and N122D. We conclude that the evolution of the R protein from a CG-sensitive ancestral state involved two epistatically-interacting substitutions (Q111R and N122D) in conjunction with compensatory effects of 10 additional substitutions that mitigate the trade-off between toxin resistance and native enzyme activity.

The adaptive division-of-labor between the R and S paralogs of ATP1A1 in *Leptodactylus* has been maintained by strong selection that has counteracted the homogenizing effects of frequent NAGC over the 35 million-year history of this genus. Similar signatures of selection to maintain sequence differentiation between neofunctionalized duplicates have been observed for the RHCE/RHD antigen proteins of humans (*35*), “major facilitator family” transporter proteins in *Drosophila* (*36*) and red/green opsins of primates (*30*). To our knowledge, only the case of opsins has been linked directly to functional differentiation, notably two closely-linked amino substitutions contributing to a red to green shift in absorbance maxima (*37*). Our study highlights similar signatures of selection not only on the two amino acid substitutions directly linked to adaptive differentiation for CG-resistance, but also at 10 more amino acid substitutions scattered throughout the protein that facilitate this neofunctionalization. Thus, by identifying interactions between adaptive substitutions and the genetic backgrounds that permit these changes, our combination of evolutionary and functional analyses reveals how mechanisms of adaptation are shaped by intramolecular epistasis and pleiotropy.

## Acknowledgments

We thank M. Przeworski for helpful comments on the manuscript. We thank C. Natarajan, K. Rohlfing, V. Wagschal, and P. Kowalski for assistance in the laboratory.

## Author contributions

PA and AJC conceived of and oversaw the project; LY, MPRO, SHA, JP, and AJC collected samples and generated sequence data; LY, AH, PA, SHA, and KZ performed evolutionary and population genetics analyses; SM, JFS, SD, AJC and PA designed functional experiments; SM and PA performed experiments and statistical analyses; SM, JFS, LY, AH and PA wrote the paper; All authors edited the manuscript;

## Data and materials availability

BioProject PRJNA627222, SRA: SRR11583961-91, nucleotides: MT396181-92, genes: MT422192-MT422203.

## Supplementary Materials

Materials and Methods

Figures S1-S7

Tables S1-S8

References 38-67

