## Supplemental Materials for "Concerted evolution reveals co-adapted amino acid substitutions in Na^+^K^+^-ATPase of frogs that prey on toxic toads"

**This PDF file includes:**

Table of Contents
Materials and Methods
Tables S1 to S8
Figs. S1 to S7

### Table of Contents

#### Materials and Methods


Sample collection and data sources 5

RNA-seq based gene discovery of ATP1A paralogs 5

Targeted sequencing of protein-coding regions of ATP1A1 paralogs 6

*De novo* genome assembly of *Leptodactylus fuscus* 6

Targeted long-read sequencing of intronic sequences of ATP1A1 7

Estimation of genealogical relationships 8

Maximum likelihood analysis of site-wise support for alternative tree topologies 9

Theoretical single-site model for the probability of maintaining an adapted substitution 10

Simulations of ATP1A1 gene family evolution 13

Inference of evolutionary parameters using Approximate Bayesian Computation 16

Construction of expression vectors 19

Generation of recombinant viruses and transfection into Sf9 cells 21

Preparation of Sf9 cell membranes 22

Verification by SDS-PAGE and western blotting 23

Ouabain inhibition assay (measurement of CG resistance) 23

ATP hydrolysis assay (measurement of ATPase activity as a proxy for protein activity) 24

Statistical analyses of biochemical assay results 24

### Supplementary Tables


Table S1. Collection information for samples of leptodactylid frogs used in this study 26

Table S2. Sources of ATP1A1 sequences included in the phylogenetic analysis 27

Table S3. Summary of the *de novo* genome assembly of *Leptodactylus fuscus* 29

Table S4. Fisher's exact test (FET) for the likelihood of site-wise support of "Non-Concerted" and "Concerted" topologies. 30

Table S5. List of primers used for intron sequencing 31

Table S6. List of engineered ATP1A1 constructs used to test functional effects of amino acid substitutions in *Leptodactylus*. 32

Table S7. Summary of the ouabain sensitivity and catalytic properties of $Na^{+}$, $K^{+}$-ATPase for each ATP1A1 construct. 33

Table S8. Statistical analysis of ouabain sensitivity and ATPase activity. 34

**Supplementary Figures**

Figure S1. Proportion of ATP1A1, ATP1A2, and ATP1A3 paralogs in brain, muscle, and stomach of seven Anuran species. 35

Figure S2. Variation among sites implicated in CG-resistance for ATPA paralogs of various species. 36

Figure S3. Positions of 12 R-copy-specific amino acid substitutions on the crystal structure of pig ATP1A1 bound to the cardiac glycoside bufalin. 37

Figure S4. Annotation of ATP1A1S and ATP1A1R paralogs in *Leptodactylus fuscus de novo* genome assembly. 38

Figure S5. A phylogenetic tree of *Leptodactylus* and outgroup species. 39

Figure S6. Western blot analysis of $Na^+$, $K^+$-ATPase with engineered ATP1A1 (α) subunits produced in this study. 40

Figure S7. Cardiac glycoside (ouabain) inhibition curves for six each recombinant *Leptodactylus* $Na^+$, $K^+$-ATPase produced in this study 41

**Materials and Methods**

Sample collection and data sources

We sampled tissues from 16 anuran species. Five *Leptodactylus* species, as well as *Engystomops pustulosus*, *Lithodytes lineatus*, and *Rhinella marina*, were collected from different geographic locations in Colombia (Table S1) and stored in RNAlater (Invitrogen) at -80ºC until used. Atissue sample of the toad, *Atelopus zeteki*, was donated by the Smithsonian's National Zoo and came from a necropsied captive animal. The outgroup species, *Megophrys nasuta*, *Kaloula pulchra*, *Rana sphenocephala*, *Rana catesbeiana*, *Dendrobates auratus*, *Melanophryniscus stelzneri*, and *Duttaphrynus melanostictus* were obtained from the pet trade under IACUC Protocol No. 2057-16. Live animals were euthanized under the supervision of a research veterinarian at Princeton University. To capture all three paralogs of ATP1A, we collected tissue samples from brain, skeletal muscle, and stomach – each of which highly expresses at least one of the three paralogs (*15*).

RNA-seq based gene discovery of ATP1A paralogs

Full-length coding sequences of ATP1A1, ATP1A2 and ATP1A3 were reconstructed for several species using RNA-seq based gene discovery. Total RNA was extracted from multiple tissues of 16 anuran species (Table S2) using TRIzol Reagents (Ambion, Life technologies) following the manufacturer's protocol. RNA-seq libraries were prepared with TruSeq RNA Library Prep Kit v2 (Illumina) and sequenced on Illumina HiSeq2500 (Genomics Core Facility, Princeton, NJ, USA) with either PE 75bp or SE 140bp (Table S2). Reads were trimmed and *de novo* assembled with Trinity v2.2.0 (*40*). ATP1A1 of *Xenopus laevis* (GenBank NM_001090595) was initially used to BLAST against the assembled transcripts of *L. latrans* to recover ATP1A1S and ATP1A1R, which were later used as queries to reconstruct ATP1A1 genes from other species. ATP1A paralogs for the rest of the species used in this study were mined from publicly available data (Table S2) following the same pipeline.

<u>Targeted sequencing of protein-coding regions of ATP1A1 paralogs</u>

Total RNA was extracted from *L. fuscus*, *L. insularum*, and *L. colombiensis* as described above and reverse-transcribed to single-strand cDNA using SuperScript III Reverse Transcriptase (Invitrogen). ATP1A1 was amplified using Phusion High-Fidelity DNA polymerase (Invitrogen) using forward primer: 5'-ATAAGTATGAGCCCGCAGCC-3' and reverse primer: 5'-CCAGGGCTGCGTCTGATTATG-3'. PCR products were cleaned with QIAquick PCR Purification Kit (Qiagen) and A-tailed with Taq Polymerase (NEB) before cloning into a pTOPO-TA vector (Invitrogen). The presence of the insert in the plasmid was confirmed by colony-PCR. Illumina-ready sequencing libraries of isolated plasmids were prepared with Tn5 transposase, charged with Illumina-ready indexed barcodes (*22*), and sequenced on Illumina MiSeq (Genomics Core Facility, Princeton, NJ, USA). *De novo* assembly of the cloned PCR products was performed with Velvet v1.2.10 (*41*) and Oases v0.2.8 (*42*). ATP1A1 paralogs were reconstructed by aligning with previously obtained ATP1A1 sequences of *L. latrans* and *L. pentadactylus*.

<u>*De novo* genome assembly of *Leptodactylus fuscus*</u>

High-molecular-weight genomic DNA was isolated from a single *Leptodactylus fuscus* individual (Table S1, JSM 205) following standard protocols. This was used to prepare a 10X-genomics Chromium library and sequenced on Illumina HiSeq X sequencer. Barcodes were removed using the Long Ranger basic v2.2.2 (https://support.10xgenomics.com/genome-exome/software/downloads/latest). Trimmed reads were used for k-mer estimation in Jellyfish (*33*, v2.2.7). The k-mer (k=21) frequency distribution was processed in GenomeScope (*43*) to estimate the genome size, heterozygosity, and percentage of repeat content. The linked-reads were assembled using the Supernova assembler (*44*) with "–accept-extreme-coverage" specified because the coverage was lower than recommended. The assembled genome is 2.42 Gb (16,530 scaffolds >=10 kb, scaffold N50 = 363 kb, Table S3) and was outputted in the psuedohap2 format (accession No. TBD). The completeness of the genome assembly was assessed using Benchmarking Universal Single-Copy Orthologs (BUSCOs, v4.0.5 (*45*)), and 72.6% of the BUSCO Tetrapoda gene annotations (version odb10) were identified (Table S3).

Targeted long-read sequencing of intronic sequences of ATP1A1

Intron annotations were determined by blasting (blast-2.26) the protein-coding sequences of ATP1A1 S and ATP1A1 R against the *L. fuscus* genome assembly (Figure S4). For the other four *Leptodactylus* species (*L. pentadactylus*, *L. latrans*, *L. insularum*, and *L. colombiensis*) and two outgroup species (*Engystomops pustulosus* and *Lithodytes lineatus*), introns were obtained via targeted long-read sequencing using Oxford Nanopore MinION. Genomic DNA was extracted with Agencourt DNAdvance Kit (Beckman Coulter, France) and ATP1A1 was amplified using LongAmp *Taq* PCR kit (NEB) using customized species-specific barcoded primers (Table S5). PCR products were gel confirmed and isolated using QIAquick PCR Purfication kit (Qiagen). Libraries were pooled and prepared for sequencing using Ligation Sequencing Kit SQK-LSK109 (Oxford Nanopore Technologies) following the manufacturer's protocol. 72,161 reads were generated within six hours, 89% passed the filter, and the real-time read length distribution matched that shown on the gel image of the amplicons. Base-calling from raw trace data was performed using Albacore v2.3.4 (Oxford Nanopore Technologies) and sequences were demultiplexed using LAST v980 (*46*). Reads that mapped to more than one barcode were discarded. Reads were assigned to each species based on barcodes using seqtk (*47*). Only reads of the expected length ± 200 nt were used for downstream analyses. For *Leptodactylus* species with two ATP1A1 paralogs, reads were further split by perfectly matching the 111-122 region of the two copies, which exhibit 22-25% difference in nucleotide sequences. Assembly was carried out using Canu v1.8 (*48*) and 1000 reads (i.e. 1000x coverage) were randomly selected for better performance. Reconstructed sequences were identical when different sets of 1000 reads were used. Filtered reads were mapped back to the reconstructed reference with minimap2 (*49*) and polished with racon v1.3.3 (*50*). Short-read sequencing data were generated using Tn5 transposase-based Illumina sequencing (as described above) to further correct and polish the sequences. Final sequences were aligned using MUSCLE (*51*) implemented in SeaView (*52*). The boundaries between introns and exons were manually adjusted to start with GT and end with AG. Sequences are available at Genbank MT422192 - 422203 (Table S2).

Estimation of genealogical relationships

A time-tree of anuran species in Figure 1A was derived from Feng *et al*. (*38*). Amino acid substitutions at sites that are implicated in cardenolide sensitivity (*22*) are shown. The nucleotide tree and protein tree (Figure. 1B, C) of *Leptodactylus* and outgroup species were built with the exons and introns and protein sequences (Table S2), respectively. Phylogenies for ATP1A1 were estimated using a maximum likelihood method implemented in SeaView and visualized in EvolView (*53*)

We constructed a multi-locus species tree for three *Leptodactylus* species (*L. fuscus, L. pentadactylus*, *L. latrans*) and two outgroups (*Engystomops pustulosus* and *Lithodytes lineatus*) with high-confidence split time estimates specifically for use in the analyses described in sections "*Theoretical single-site model for the probability of maintaining an adapted substitution*" and "*Simulations of ATP1A1 gene family evolution*". Protein-coding genes were predicted from *de novo* transcriptome assemblies for each species using Augustus (v3.2.2) (*54*) and queried against the Tetrapoda ortholog database (odb10, https://www.orthodb.org) using BLAST (tblastn). A concatenated multi-alignment of mRNA sequences was created for 813 orthologous proteins longer than 100 amino acids that were shared among all five species. The best-fit nucleotide substitution model for each protein (i.e. each initial partition) was first determined using the "ModelFinder" function of IQ-TREE 2 (v.2.0.4; *55*) (command line: iqtree2 -s concat_813_mafft.fasta -spp partition.txt -m MFP -nt AUTO -safe --prefix concat_813_partition_MFP). Proteins with the same inferred mutation model were subsequently merged into the same partition (using "-m TESTMERGE") prior to phylogenetic inference (command line: iqtree2 -s concat_813_mafft.fasta -spp partition_MFP_best_scheme.nex -m TESTMERGE -nt AUTO  --prefix concat_813_partition_MFP_merged).

<u>Maximum likelihood analysis of site-wise support for alternative tree topologies</u>

We used site-wise likelihoods to evaluate the relative level of statistical support for two alternative tree topologies relating to the origin of R/S ATP1A1 paralogs: Model 1 ("Non-Concerted") posits a single ancient origin of a R/S duplication with no concerted evolution:

((Lfus_S,(Lpen_S,(Lins_S,Llat_S,Lcol_S))),(Lfus_R,(Lpen_R,(Lins_R,Llat_R,Lcol_R)))). Model 2 (“Concerted”) is the expected topology under concerted evolution: ((Lfus_S, Lfus_R), ((Lpen_S, Lpen_R), ((Lins_S, Lins_R), (Llat_S, Llat_R), (Lcol_S, Lcol_R)))). We note that the speciation events are assumed to follow the order inferred in the section “*Estimation of genealogical relationships*”. For each nucleotide states (e.g. AAAATTTTTT, in the order of Lfus_S, Lfus_R, Lpen_S, LpenR, Llat_S, LlatR, LcolS, LcolR, LinsS, LinsR), likelihoods for the two topologies were calculated using *PAML 4.8 baseml* (*56*). We consider |∆log-likelihood| ≥ 2, as significant support for one topology over the other. 4-, 2-, 0-fold degenerate sites were classified using *MEGA* 7 (*57*) and all variants at these sites were categorized as either synonymous or nonsynonymous. We used Fisher’s Exact Test to test the hypothesis that the ratio of synonymous and nonsynonymous variants is independent of support for one of the topologies over the other (Table S4).

We further tested whether synonymous variants supporting alternative tree topologies (as outlined above) are equally distant from R/S distinguishing substitutions: We computed the distance of each variant from the nearest R/S distinguishing substitution, and compared the median distance of synonymous variants with |∆log-likelihood| ≥ 2 support for the “Non-Concerted” genealogy to a random sample of synonymous variants supporting multiple origins.

<u>Theoretical single-site model for the probability of maintaining an adapted substitution</u>

Below, we describe the model and parameters used to compute the probability of maintaining a diverged substitution in two gene copies.

**Model.** We consider a single biallelic amino acid site in tandemly duplicated genes, evolving for *t* years. The two gene copies are initially fixed for the two distinct alleles. The site experiences mutation at rate $2\mu$ (or $4\mu$ for both copies) where $\mu$ is the per-nucleotide mutation rate, assuming for simplicity that all sites are biallelic, all mutations in the first two positions of the codon are nonsynonymous and all mutations at the third position are synonymous. The site also experiences non-allelic gene conversion at rate $4c$ (for both copies) and is under purifying selection with fitness cost $s > 0$, such that having two distinct alleles at the two copies confers a fitness of 1 and having the same allele confers to fitness $(1 - s)$.

*De novo* mutations (through point mutation or gene conversion) from the initial distinct-allele haplotype to a same-allele haplotype can occur in all haplotypes in the population. In a diploid population of size $N$, de novo same-allele haplotypes arise at rate

$$P(de\ novo\ same - allele\ haplotype) = \ 2N \cdot 4 \cdot (\mu + c).$$

The probability of fixation is bounded by the neutral case of $s = 0$, such that

$$P(same - allele\ haplotype\ fixes) < \frac{1}{2N}.$$

If

$$8N \cdot (\mu + c) \ll 1$$

and

$$\frac{1}{2N} \ll 1,$$

then the overall per-year rate of fixation for deleterious haplotypes, $\alpha$, can be approximated by the product of these two,

$$\alpha = \ P(de\ novo\ same - allele\ haplotype) \cdot \ P(same - allele\ haplotype\ fixes) =$$

$$8N(c + \mu) \cdot \frac{e^{s} - 1}{e^{2Ns} - 1},$$

where we replaced $P(deleterious\ haplotype\ fixes)$ with Kimura’s fixation probability for a deleterious allele (*58*, *59*). Assuming a vanishingly small probability of back-mutations—

namely, that no fixation of a same-allele haplotype is followed by another fixation reversing the haplotype back to the distinct alleles—the probability of maintaining the distinct-alleles haplotype for $t$ years is:

$$P(maintenance\ of\ distinct\ alleles) = (1-\alpha)^t = \left(1 - 8N(c+\mu)\frac{e^s - 1}{e^{2Ns} - 1}\right)^t. \quad (1)$$

We note that if $s \ll 1$ then

$$e^s \approx 1 + s,$$

and therefore

$$P(maintenance\ of\ distinct\ alleles) \approx \left(1 - 4(c+\mu)\frac{2Ns}{e^{2Ns} - 1}\right)^t, \quad (2)$$

giving a maintenance probability that is only dependent on the effective population size and the selection coefficient through the compound population parameter $2Ns$.

**Parameters.** To compute maintenance probabilities, we set the point mutation rate to its estimate by Sun *et al.* (*60*) (also supported by earlier work from Crawford (*61*)) of

$$\mu = 0.776 \cdot 10^{-9} \text{ mutations per bp per year.} \quad (3)$$

We wished to use the total branch length of the *Leptodactylus* phylogeny for $t$, the maintenance time, to reflect the observation of trans-specific maintenance. In considering the phylogenetic tree and split times here and in the evolutionary simulations of the section "Simulations of ATP1A1 gene family evolution" below, we only considered a subset of three *Leptodactylus* species— *L. fuscus, L. latrans* and *L. pentadactylus*—for which confident species split time estimates were available (see "*Estimation of genealogical relationships*" section; Fig. S5): a split between *L. fuscus* and the common ancestor of the two other species 29,187,798 years ago, followed by a split between *L. latrans* and *L. pentadactylus* 27,426,120 years ago. Therefore, the total time on the species tree was set to

$$t = 2 \cdot 29{,}187{,}798 + 27{,}426{,}120 = 85{,}801{,}716\, years. \quad (4)$$

The maintenance probabilities shown in Fig. 3A were computed using eq. (1), plugging in the parameters in eq. (3) and (4) and across a grid of $(Ns) \in [-1,1.5]$ and $c \in [0,2.5]$ values.

Simulations of ATP1A1 gene family evolution

**Overview.** We developed evolutionary simulations with the goal of gauging the evolutionary parameters that could have produced the observed spatial divergence patterns along ATP1A1. Typically, and whenever possible, analytic likelihood or posterior probability functions are derived for such a task. Alternatively, backward-in-time simulations are used, because of their high computational efficiency. However, analytic or backward-in-time approaches were intractable for our purposes: both because we wished to account for the spatial divergence patterns and not consider sites independently—and because our model of ATP1A1 evolution in *Leptodactylus* includes complex interactions between point mutation, NAGC, and selection that violate typical assumptions of analytic / backward in time sequence evolution models. We therefore developed a forward-in-time simulation of R and S. The simulations take a set of parameters $\Theta$ as input (see section "Fitness model and other parameterization" below), start with two ancestral sequences and end with an output of contemporary R and S sequences in multiple *Leptodactylus* species, which we later compare to the observed data (see section "Inference of evolutionary parameters using Approximate Bayesian Computation").

**Fitness model and other parameterization.** At the heart of our simulation, we consider the possible fixation of new haplotypes in *Leptodactylus* lineages. These fixations follow random occurrence of de novo point mutations or NAGC in one of the haplotypes in the population; but the probability of fixation on the lineage will depend on the selection acting on the novel variant.

The ancestral haplotype with which the simulation begins is assumed to underlie the optimal function of R, S and interactions between them, and thus to be of optimal fitness. Therefore, the absolute fitness $f$ of a haplotype $X$ at any point of the simulation depends on its divergence from the ancestral haplotype with which the simulation begins, as follows:

$$f(X) = s_1X_1 + s_2X_2 + s_yY + s_zZ + s_{12}X_1X_2 + s_{1y}X_1Y + s_{2y}X_2Y,$$

where $X_1 \in \{0,1,2\}$ is the number of residue differences between $X$ and the ancestral haplotype at position 111 of the amino acid sequences of both R and S; $X_2 \in \{0,1,2\}$ is the number of residue differences between $X$ and the ancestral haplotype at position 122; $Y \in \{0,1,\ldots,20\}$ is the number of residue differences between $X$ and the ancestral haplotype at the other 10 R/S distinguishing substitutions (referring to the substitutions strongly distinguishing R and S in the observed sequences); and $Z$ is the number of total residue differences between $X$ and the ancestral haplotype in the rest of the amino acid sequence. $\{s_1, s_2, s_y, s_z, s_{12}, s_{1y}, s_{2y}\}$ represent selection coefficients and are fixed parameters that are taken as input of the simulation.

Other parameters taken as input by our simulation (see pseudocode below) include:

- $N$, the population size of each extant *Leptodactylus* lineage
- $\mu$, the per haplotype, per nucleotide per year mutation rate.
- $l$, the mean NAGC tract length in base pairs. We model the tract length as Geometrically distributed (*39*, *62*).
- $c$, the NAGC per nucleotide per year rate. Note that this is the rate in which a site is included in a NAGC tract, not the rate at which NAGC events initiate at the site.
- A rooted species tree, consisting of a bifurcating topology and branch lengths (split times) in years.

**Simulation pseudocode:**

1. *Initialize time t to the TMRCA of all species.*
2. *While t < today,*

*2.1. Advance t by* $t_w$*, the waiting time for the next mutational event, where*

$$t_w \sim Exp((2N\ haplotypes) \cdot \big(extant\ species\big) \cdot (2\ paralogs\ per\ species) \cdot (ATP1A1\ sequence\ length) \cdot (rate\ per\ nucleotide\ c + \mu))$$

*2.2 If t > time for lineage split that had not yet occurred,*

*2.2.1 bifurcate lineage: copy R and S sequences of ancestral lineage into an identical copy and label each of the two sets as one of the lineages.*

*2.3 Draw* $U_{event} \sim U(0,1)$. *if* $U_{event} < \frac{\mu}{\mu + c}$ *then the de novo mutational event is a point mutation, else, it is a NAGC event.*

*2.4 Draw (uniformly) an extant species in which the event occurred.*

*2.5 Draw (uniformly) a paralog (R or S) in which the mutation occurred or served as the template for NAGC.*

*2.6 Draw (uniformly) a random nucleotide position where the mutational event occurred.*

*2.7 If the de novo event is a NAGC event,*

*2.5.1 Draw a tract length* $L \sim Geo(l)$. *Expand tract around initiation site, with a uniform fraction extending to the left and right of the site.*

*2.8 Translate the derived, de novo haplotype and the ancestral haplotype to amino acid sequences and calculate their fitness; calculate the resulting relative fitness of the derived haplotype.*

*2.9 Calculate $p_{fix}$, the fixation probability (see below) for a haplotype at frequency $\frac{1}{2N}$ conferring relative fitness as calculated in 2.8.*

*2.10 Draw $U_{fix} \sim U(0,1)$. If $U_{fix} < p_{fix}$,*

*2.10.1 Fix: Replace ancestral haplotype in the species with the de novo haplotype.*

In step *2.9*, we consider a *de novo* haplotype arising in the population (namely, at frequency $\frac{1}{2N}$) with relative fitness $1 + s$ to have probability

frequency $\frac{1}{2N}$) with relative fitness $1 + s$ to have probability

$$p_{fix} = \begin{cases} \frac{e^{s} - 1}{e^{2Ns} - 1} & if\ s < 0\ (deleterious) \\ \frac{1}{2N} & if\ s = 0\ (neutral) \\ \frac{1 - e^{-s}}{1 - e^{-2Ns}} & if\ s > 0\ (advantageous) \end{cases}$$

of fixing in the population, following Kimura (*58*).

of fixing in the population, following Kimura (*58*).

Inference of evolutionary parameters using Approximate Bayesian Computation

**Overview.** We used an Approximate Bayesian Computation (ABC) approach to estimate evolutionary parameters, including gene conversion rates and the strength of purifying selection acting at different sites in ATP1A1. In each iteration $j$, we sampled a set of parameters $\Theta_j$ from a predefined prior distribution. We approximated the posterior distribution of $\Theta_j$ by the empirical distribution given by a subset of this sample that generates divergence patterns that we inferred as closest to the true data. To infer the "distance" of simulated data from the observed data, we ran forward-in-time evolutionary simulations of ATP1A1 sequence evolution and quantified the similarity of the simulated divergence patterns to the observed divergence patterns. Simulations all begin with the same ancestral R and S genes in a common ancestor, and end with six evolved (simulated) contemporary sequences, corresponding to R and S in three *Leptodacylus* species. From the divergence patterns between these six simulated sequences, we computed $d(\Theta_j)$, the distance between the simulated and the observed (real sequence data) ATP1A1 divergence patterns.

**Parameter set and prior distribution.** Our evolutionary simulations take as input a set of parameters as defined in the section "Simulations of ATP1A1 gene family evolution",

$$\Theta = \{\mu, c, l, N, s_1, s_2, s_z, s_y, s_{12}, s_{1y}, s_{2y}\}.$$

The prior distributions of single parameters are mutually independent. Namely, the prior distribution on $\Theta$ was set as

$$\pi(\Theta) = \pi_c(c)\pi_{\tilde{s}}(\tilde{s})\pi_{s_z}(s_z),$$

where $\pi_K$ is the marginal prior distribution of $K$, and $\tilde{s} \coloneqq s_1 = s_2 = s_y$ such that all 12 sites distinguishing R and S in the observed data are under the same selective constraint, but it is free to differ from the selective constraint on other amino acids. The marginal priors on the gene conversion rate $c$ and selection coefficients $\tilde{s}$, $s_z$ were set as

$$\log_{10}(\frac{c}{\mu}) \sim U(0{,}2.5),$$

$$\log_{10}(N\tilde{s}) \sim U(-1{,}1)$$

and

$$\log_{10}(Ns_y) \sim U(-1{,}1).$$

The other parameters were assumed fixed: we set the mutation rate to be $\mu = 0.776 \cdot 10^{-9}$ mutations per bp per year and the diploid population size (in each extant species at a given time in the simulation) to be N=10 (2N=20) as in the section "Theoretical single-site model for the probability of maintaining an adapted substitution". This small population size was chosen to allow for computational efficiency, because the simulation run time scaled linearly with N, and our inference became computationally infeasible with substantially larger population sizes. The mean tract length for gene conversion events was set to $l = 100bp$. We have found that there is very limited resolution given by our inference scheme on selection interaction terms, $s_{12}, s_{1y}$ and $s_{2y}$ when we allowed them to vary. We therefore set these fitness interaction terms to zero.

**Measuring similarity to observed divergence patterns.** Given $y$, a set of R and S nucleotide sequences in three species, we computed two summaries of the divergence at each nucleotide site $i$: $d_o(y_i)$, the sum of pairwise Hamming distances between R sequences in a pair of species (each $\in \{0,1\}$ since only one site is considered) plus the sum of pairwise Hamming distances between R sequences; and $d_p(y_i)$, the sum—across the three species—of Hamming distances between paralogous R and S sequences. Let $y^{obs}$ be the six observed sequences and $y^{\Theta j}$ be the sequences output at the end of simulation run j. We measured the divergence between the simulated and observed data at site I as

$$d_i(\Theta_j) = d_i(y^{obs}, y^{\Theta j}) = d_o\left(y_i^{obs}, y_i^{\Theta j}\right) + d_p\left(y_i^{obs}, y_i^{\Theta j}\right).$$

This per-site distance was computed for all positions $I$, namely nucleotide sites without missing data or insertions/deletions in any of the six observed sequences. Finally, the distance between simulation $j$ and the observed data is given by

$$d(\Theta_j) = \sum_{\text{sites i}} w_i d_i(y^{obs}, y^{\Theta j}),$$

where $w_i$ are position-importance weights, giving extra weight for divergence patterns near R/S distinguishing sites. These weights were set as

$$w_i = 1 + \sum_{k=1}^{12} 10 \cdot e^{-|i-i_k|},$$

where $\{i_k\}$ is the set of $12 \cdot 3$ positions coding for one of the 12 R/S distinguishing substitution sites.

**Analysis.** We ran 23,323 simulations with Θ sampled from its prior distribution. We kept ~1% of these parameter sets—234 sets which produced simulations with the lowest $d(\cdot)$ values, and considered them as samples from the approximate posterior distribution. We then used the functions *kde3d* (for the approximate posterior distribution of $c, s_z$ and $\tilde{s}$) and *kde2d* (for the marginal approximate posterior distribution of $c$ and $\tilde{s}$) from the *R* packages *misc3d* (*63*) and *MASS* (*64*) to estimate the posterior with a spline fit using over 200 bins per dimension, in the range set by our prior distribution on each parameter, and with otherwise default settings of *kde3d and kde2d*. The approximate posterior mode was

$$(c = 18\mu, 2N\tilde{s} = 6, 2Ns_z = 1),$$

and the marginal posterior mode on the first two parameters was

$$(c = 9\mu, 2N\tilde{s} = 7).$$

The (single dimension) marginal credible interval mentioned in the main text are high posterior density credible intervals.

<u>Construction of expression vectors</u>

$Na^+$,$K^+$-ATPase is a multi-subunit protein that requires co-expression of the alpha (ATP1A) and beta subunits (ATP1B) in cell lines. An RNA-seq analysis of *Leptodactylus* brain, stomach, and muscle tissues revealed that ATP1B1, one of four paralogous copies of ATP1B, is the most ubiquitously expressed. cDNA was reverse transcribed from *Leptodactylus latrans* stomach mRNA using the Superscript III Reverse Transcriptase kit (Invitrogen™). The ATP1B1 gene was amplified from cDNA with the primers, 5’ATCCTCGAGATGGCCAGAGACAAAACCAAGGA 3’ and 5’ TGTGGTACCTCAGCTACTCTTAATCTCCAACTTTA 3’, which added a XhoI site at the 5’ end and a KpnI site at the 3’ end. ATP1B1 amplicons were inserted into pFastBac Dual expression vectors (Life Technologies) at the p10 promoter with XhoI and KpnI (FastDigest; Thermo Scientific™), and then control sequenced. The vector insert sequence was an identical match to the *L. latrans* $\beta$1-subunit transcript generated in this study. ATP1A1S was amplified from cDNA with the primers 5’ TAATACTAGTATGGGATACGGGGCCGGACGTGAT 3’ and 5’ ACTGCGGCCGCTTAATAATAGGTTTCTTTCTCCA 3’ and ATP1A1R was amplified from a previously constructed vector containing a truncated copy of the gene with the overhang primers 5’ TAATACTAGTATGGGATACGGGGCCGGACGTGATGAGTATGAGCCCGCAGCCACTTCTGAACATGGCGGCAAGAAGAAAGGCAAAGGGAAGGATAAGGAT 3’ and 5’ ACTGCGGCCGCTTAATAATAGGTTTCTTTCTCCACCCAGCCGCCAGGGCTGCGTCTGATTATCAGTTTTCGGATTTCATCATATATGAAGATGAGCAGAGAGTAGGGGAAGGCACAGAACCACCATGTTGGTTTCAGTGGGTACATGCGGAGTGCCACATCCATGCCTGGG 3’. Both pairs of primers added a SpeI site at the 5’ end and a NotI site at the 3’ ends. All gene amplifications were performed using a high-fidelity proofreading polymerase (Phusion High-Fidelity DNA Polymerase; Thermo Scientific™). ATP1A1S and ATP1A1R amplicons were inserted at the $P_{PH}$ promoter of pFastBac Dual expression vectors already containing ATP1B1 with SpeI and NotI (FastDigest; Thermo Scientific™), and then control sequenced. The ATP1A1S sequence was an identical match to the *L. latrans* sensitive α1-subunit transcripts and the ATP1A1R sequence was an identical match to *L. latrans* resistant α1-subunit transcripts generated from this study. *Escherichia coli* DH5α cells (Invitrogen™) were transformed with the two resulting expression vectors (pFastBac Dual + ATP1B1 + ATP1A1S and pFastBac Dual + ATP1B1 + ATP1A1R). These completed vectors were then used to introduce the amino acid codons of interest by site-directed mutagenesis (QuikChange II XL Kit; Agilent Technologies, La Jolla, CA, USA) according to the manufacturer's protocol. One ATP1A1S gene construct was synthesized by Invitrogen™ GeneArt (S+12R). All resulting vectors had the α1-subunit gene under the control of the $P_{PH}$ promoter and the $\beta$1-subunit gene under the p10 promoter (Table S6).

Generation of recombinant viruses and transfection into Sf9 cells

*Escherichia coli* DH10bac cells harboring the baculovirus genome (bacmid) and a transposition helper vector (Life Technologies) were transformed according to the manufacturer's protocol with expression vectors containing the different gene constructs. Recombinant bacmids were selected through PCR screening, grown, and isolated (*65*). Subsequently, Sf9 cells ($4 \times 10^5$ cells*ml) in 2 ml of Insect-Xpress medium (Lonza, Walkersville, MD, USA) were transfected with recombinant bacmids using Cellfectin reagent (Life Technologies). After a three-day incubation period, recombinant baculoviruses were isolated (P1) and used to infect fresh Sf9 cells ($1.2 \times 10^6$ cells*ml) in 10 ml of Insect-Xpress medium (Lonza, Walkersville, MD, USA) with 15 mg/ml gentamycin (Roth, Karlsruhe, Germany) at a multiplicity of infection of 0.1. Five days after infection, the amplified viruses were harvested (P2 stock).

Preparation of Sf9 cell membranes

For production of recombinant $Na^+$,$K^+$-ATPase, Sf9 cells were infected with the P2 viral stock at a multiplicity of infection of 1e3. The cells (1.6 x $10^6$ cells*ml) were grown in 50 ml of Insect-Xpress medium (Lonza, Walkersville, MD, USA) with 15 mg/ml gentamycin (Roth, Karlsruhe, Germany) at 27°C in 500 ml flasks (*33*). After 3 days, Sf9 cells were harvested by centrifugation at 20,000 x g for 10 min. The cells were stored at -80 °C, and then resuspended at 0 °C in 15 ml of homogenization buffer (0.25 M sucrose, 2 mM EDTA, and 25 mM HEPES/Tris; pH 7.0). The resuspended cells were sonicated at 60 W (Bandelin Electronic Company, Berlin, Germany) for three 45 s intervals at 0 °C. The cell suspension was then subjected to centrifugation for 30 min at 10,000 x g (J2-21 centrifuge, Beckmann-Coulter, Krefeld, Germany). The supernatant was collected and further centrifuged for 60 min at 100,000 x g at 4 °C (Ultra- Centrifuge L-80, Beckmann-Coulter) to pellet the cell membranes. The pelleted membranes were washed once and resuspended in ROTIPURAN® p.a., ACS water (Roth) and stored at -20 °C. Protein concentrations were determined by Bradford assays using bovine serum albumin as a standard. Six biological replicates were produced for each $Na^+$,$K^+$-ATPase construct.

Verification by SDS-PAGE and western blotting

For each biological replicate, 50 ug of protein were solubilized in 4x SDS-polyacrylamide gel electrophoresis sample buffer and separated on SDS gels containing 10% acrylamide. Subsequently, they were blotted on nitrocellulose membrane (HP42.1, Roth). To block non-specific binding sites after blotting, the membrane was incubated with 5% dried milk in TBS-Tween 20 for 1 h. After blocking, the membranes were incubated overnight at 4 °C with the primary monoclonal antibody α5 (Developmental Studies Hybridoma Bank, University of Iowa, Iowa City, IA, USA). Since only membrane proteins were isolated from transfected cells, detection of the α subunit also indicates the presence of the β subunit. The primary antibody was detected using a goat-anti-mouse secondary antibody conjugated with horseradish peroxidase (Dianova, Hamburg, Germany). The staining of the precipitated polypeptide-antibody complexes was performed by addition of 60 mg 4-chloro-1 naphtol (Sigma-Aldrich, Taufkirchen, Germany) in 20 ml ice-cold methanol to 100 ml phosphate buffered saline (PBS) containing 60 ul 30% $H_2O_2$. See Fig. S6.

Ouabain inhibition assay (measurement of CG resistance)

To determine the sensitivity of each $Na^+$,$K^+$-ATPase construct against the water-soluble cardiac glycoside, ouabain (Acrōs Organics), 100 ug of each protein was pipetted into each well in a nine-well row on a 96-well microplate (Fisherbrand) containing stabilizing buffers (see buffer formulas in (*66*). Each well in the nine-well row was exposed to exponentially decreasing concentrations ($10^{-3}$ M, $10^{-4}$ M, $10^{-5}$ M, $10^{-6}$ M, $10^{-7}$ M, $10^{-8}$ M, dissolved in distilled $H_2O$) of ouabain, distilled water only (experimental control), and a combination of an inhibition buffer lacking KCl and $10^{-2}$ M ouabain to measure background ATPase activity (see (*66*)). The proteins were incubated at 37°C and 200 rpms for 10 minutes on a microplate shaker (Quantifoil Instruments, Jena, Germany). Next, ATP (Sigma Aldrich) was added to each well and the proteins were incubated again at 37°C and 200 rpms for 20 minutes. The activity of $Na^+$,$K^+$-ATPases following ouabain exposure was determined by quantification of inorganic phosphate (Pi) released from enzymatically hydrolyzed ATP. Reaction Pi levels were measured according to the procedure described by Taussky and Shorr (*67*) (see (*66*)). All assays were run in duplicate and the average of the two technical replicates was used for subsequent statistical analyses. Absorbance for each well was measured at 650 nm with a plate absorbance reader (BioRad Model 680 spectrophotometer and software package).

ATP hydrolysis assay (measurement of ATPase activity as a proxy for protein activity)

To determine the functional efficiency of different $Na^+$,$K^+$-ATPase constructs, we calculated the amount of Pi hydrolyzed from ATP per mg of protein per minute. The measurements were obtained from the same assay as described above. In brief, absorbance from the experimental control reactions, in which 100 ug of protein was incubated without any inhibiting factors (i.e., ouabain or buffer excluding KCl), were measured and translated to mM Pi from a standard curve that was run in parallel (1.2 mM Pi, 1 mM Pi, 0.8 mM Pi, 0.6 mM Pi, 0.4 mM Pi, 0.2 mM Pi, 0 mM Pi).

Statistical analyses of biochemical assay results

Background phosphate absorbance levels from reactions with inhibiting factors were used to calibrate phosphate absorbance in wells measuring ouabain inhibition and in the control wells (*66*). For ouabain sensitivity measurements, calibrated absorbance values were converted to percentage non-inhibited $Na^+$,$K^+$-ATPases activity based on measurements from the control wells (*66*). These data were plotted and log $IC_{50}$ values were obtained for each biological replicate from nonlinear fitting using a four-parameter logistic curve, with the top asymptote set to 100 and the bottom asymptote set to zero (Fig. S7). Curve fitting was performed with the nlsLM function of the minipack.lm library in R. For comparisons of recombinant protein ATPase activity, the calculated Pi concentrations of 100 ug of protein assayed in the absence of ouabain were converted to nmol Pi/mg protein/min. We used ANOVA to test for effects of substitutions on ouabain resistance (log $IC_{50}$) and enzyme activity (Table S8; Levene's Test for Homogeneity of Variance for $IC_{50}$: $F_{7,40}$=0.68 p=0.69 and enzyme activity: $F_{7,40}$=0.31 p=0.94). We used linear regression to estimate effect sizes associated with substitutions and pairwise t-tests to identify significant differences between substitution combinations (Table S8). All statistical analyses were implemented in R.

**Table S1. Collection information for samples of leptodactylid frogs used in this study.** ANDES-A refers to the Amphibian collection of the *Museo de Historia Natural C. J. Marinkelle* of the Universidad de los Andes, Bogotá, Colombia. Collector acronyms are Andrew J. Crawford (AJC), Juan Salvador Mendoza (JSM), Juan Manuel Padial (JMP). All collecting sites are located in Colombia (CO). Samples without museum voucher IDs are in the process of being accessioned into the ANDES-A collection.

| Species | Museum ID | Field ID | Data type | Locality | Latitude, longitude |
|---|---|---|---|---|---|
| *Engystomops pustulosus* | | AJC 3734 | RNA-seq | Mariquita, Tolima, CO. | 05.2635, -074.891 |
| *Engystomops pustulosus* | | JSM 228 | intron | Zambrano, Bolívar, CO | 09.75, -074.8333 |
| *Lithodytes lineatus* | | AJC 6408 | RNA-seq | *pending* | |
| *Lithodytes lineatus* | ANDES-A 2536 | AJC 2406 | intron | Trubon, Río Vaupés, Vaupés, CO. | 01.21, -070.62 |
| *Leptodactylus fuscus* | ANDES-A 3141 | AJC 5344 | plasmid | Neiva, Huila, CO. | 02.8796, -075.2757 |
| *Leptodactylus fuscus* | | JSM 205 | genome | Garzón, Huila, CO. | 02.2058, -075.6440 |
| *Leptodactylus pentadactylus* | ANDES-A 2327 | AJC 4761 | RNA-seq | Leticia, Amazonas, CO. | -03.865, -070.2061 |
| *Leptodactylus pentadactylus* | ANDES-A 949 | JMP 2179 | intron | Leticia, Amazonas, CO. | -04.10592, -069.25 |
| *Leptodactylus latrans* | | AJC 3653 | RNA-seq | Puerto Carreño, Vichada, CO. | 06.10, -067.483 |
| *Leptodactylus latrans* | ANDES-A 1148 | AJC 3430 | intron | Orocué, Casanare, CO. | 04.9093, -071.4286 |
| *Leptodactylus insularum* | ANDES-A 3146 | AJC 5345 | | Neiva, Huila, CO. | 02.8441, -075.3328 |
| *Leptodactylus insularum* | | AJC 3752 | CDS | Montería, Córdoba, CO. | 08.7917, -075.8629 |
| *Leptodactylus insularum* | | JSM 261 | intron | Barú, Bolívar, CO. | 10.1458, -075.6792 |
| *Leptodactylus colombiensis* | | AJC 5510 | | Santa María, Boyacá, CO. | 04.8499, -073.2653 |
| *Leptodactylus colombiensis* | ANDES-A 3066 | AJC 3755 | CDS | Nilo, Cundinamarca, CO. | 04.3584, -074.5649 |
| *Leptodactylus colombiensis* | | AJC 4301 | intron | San Martín, Meta, CO. | 03.6969, -073.6986 |

**Table S2. Sources of ATP1A1 sequences included in the phylogenetic analysis (Fig. 1)**. New data generated by this study are indicated by blue text (RNA-seq datasets: PRJNA627222, genome assembly: PRJNA631731).

| Species | Data type and format | Accession |
|---|---|---|
| *Atelopus zeteki* | RNA-seq, PE 140 bp | skin: SRR11583991 |
| *Bombina maxima* | RNA-seq, PE 90 bp | skin: SRR566619 |
| *Bufotes viridis* | RNA-seq, PE 100 bp | SRR2163277 |
| *Craugastor fitzingeri* | RNA-seq, SE 100 bp | skin: SRR1560905 |
| *Cyclorana alboguttata* | RNA-seq, SE 105 bp | muscle: SRR619475 |
| *Dendrobates auratus* | RNA-seq, PE 150 bp | brain: SRR11583990<br>muscle: SRR11583979<br>stomach: SRR11583968 |
| | *de novo* assembly | CDS: MT813444 |
| *Duttaphrynus melanostictus* | RNA-seq, PE 150 bp | brain: SRR11583966<br>muscle: SRR11583965<br>stomach: SRR11583964 |
| | *de novo* assembly | CDS: MT813445 |
| *Engystomops pustulosus* | RNA-seq, PE 140 bp | brain: SRR11583963<br>stomach: SRR11583962 |
| | *de novo* assembly | CDS: MT396181 |
| | long-read sequencing | partial gene: MT422192 |
| *Fejervarya cancrivora* | RNA-seq, PE 100 bp | SRR1554290 |
| *Homo sapiens* | NCBI reference sequence | NM_001160233.1 |
| *Kaloula pulchra* | RNA-seq, PE 150 bp | muscle: SRR11583961<br>stomach: SRR11583989 |
| | *de novo* assembly | CDS: MT813446 |
| *Leptobrachium boringii* | RNA-seq, PE 100 bp | SRR4436787 |
| *Leptodactylus colombiensis* | cloning, plasmid sequencing | CDS: MT396187 (ATP1A1S)<br>MT396188 (ATP1A1R) |
| | long-read sequencing | partial gene: MT422198 (ATP1A1S)<br>MT422199 (ATP1A1R) |
| *Leptodactylus fuscus* | cloning, plasmid sequencing | CDS: MT396183 (ATP1A1S)<br>MT396184 (ATP1A1R) |
| | single-molecule genomic sequencing | *de novo* assembly: TBD<br>partial gene: MT422194 (ATP1A1S)<br>MT422195 (ATP1A1R) |
| *Leptodactylus insularum* | cloning, plasmid sequencing | CDS: MT396191 (ATP1A1S)<br>MT396192 (ATP1A1R) |
| | long-read sequencing | partial gene: MT422202 (ATP1A1S)<br>MT422203 (ATP1A1R) |
| *Leptodactylus latrans* | RNA-seq, SE 140 bp | brain: SRR11583988<br>stomach: SRR11583987 |
| | *de novo* assembly | CDS: MT396189 (ATP1A1S)<br>MT396190 (ATP1A1R) |
| | long-read sequencing | partial gene: MT422200 (ATP1A1S)<br>MT422201 (ATP1A1R) |
| *Leptodactylus pentadactylus* | RNA-seq, SE 140 bp | brain: SRR11583986<br>stomach: SRR11583985 |

| | | |
|---|---|---|
| | *de novo* assembly | CDS: MT396185 (ATP1A1S)<br>MT396186 (ATP1A1R) |
| | long-read sequencing | partial gene: MT422196 (ATP1A1S)<br>MT4221957 (ATP1A1R) |
| *Limnodynastes peronii* | RNA-seq, PE 100 bp | SRR8712702 |
| *Lithodytes lineatus* | RNA-seq, PE 75 bp | muscle: SRR11583984<br>stomach: SRR11583983 |
| | *de novo* assembly | CDS: MT396182 |
| | long-read sequencing | partial gene: MT422193 |
| *Mantella betsileo* | RNA-seq, PE 90 bp | skin: SRR7592160 |
| *Megophrys nasuta* | RNA-seq, PE 150 bp | brain: SRR11583982<br>muscle: SRR11583981<br>stomach: SRR11583980 |
| | *de novo* assembly | CDS: MT813448 |
| *Melanophryniscus stelzneri* | RNA-seq, PE 150 bp | brain: SRR11583978<br>muscle: SRR11583977<br>stomach: SRR11583976 |
| | *de novo* assembly | CDS: MT813449 |
| *Odorrana tormota* | RNA-seq, PE 150 bp | skin: SRR6896138 |
| *Oreobates cruralis* | RNA-seq, PE 126 bp | intestine: SRR5507183 |
| *Oreolalax rhodostigmatus* | RNA-seq, PE 150 bp | SRR6265740 |
| *Pelobates fuscus* | RNA-seq, PE 90 bp | SRR5119616 |
| *Pelophylax lessonae* | RNA-seq, PE 90 bp, PE | SRR1164893 |
| *Quasipaa boulengeri* | RNA-seq, PE 100 bp, PE | SRR2962603 |
| *Rana catesbeiana* | RNA-seq, PE 150 bp | brain: SRR11583975<br>muscle: SRR11583974<br>stomach: SRR11583973 |
| | *de novo* assembly | CDS: MT813450 |
| *Rana sphenocephala* | RNA-seq, PE 150 bp | brain: SRR11583972<br>muscle: SRR11583971<br>stomach: SRR11583970 |
| | *de novo* assembly | CDS: MT813451 |
| *Rattus norvegicus* | NCBI reference sequence | NM_012504.1 |
| *Rhinella marina* | RNA-seq, PE 140 bp | brain: SRR11583969<br>skin: SRR11583967 |
| | *de novo* assembly | CDS: MT813452 |
| *Xenopus tropicalis* | NCBI reference sequence | NM_204076.1 |

**Table S3. Summary of the *de novo* genome assembly of *Leptodactylus fuscus***

| Sequencer | HiSeq X |
|---|---|
| Assembly software | Supernova 2.1.1 |
| Number of reads | 775.95 million |
| Read format | Paired end 150 bp |
| Effective read depth coverage | 48.35 |
| Estimated genome size | 2.42 Gb |
| Weighted mean molecule size | 29.36 kb |
| Number of scaffolds >= 10 kb (long scaffolds) | 16,530 |
| N50 contig size | 19.69 kb |
| N50 scaffold size | 362.61 kb |
| Assembly size (only scaffolds >= 10 kb) | 1.26 Gb |
| BUSCO version | 4.0.5 |
| Lineage dataset | Tetrapoda_odb10 |
| Input genome format | supernove pseudohap2_2 |
| Total groups searched | 5310 |
| Complete BUSCOs | 3182 (60.0%) |
| Complete and single-copy BUSCOs | 3041 (57.3%) |
| Complete and duplicated BUSCOs | 141 (2.7%) |
| Fragmented BUSCOs | 669 (12.6%) |
| Missing BUSCOs | 1459 (27.4%) |

**Table S4. Fisher's exact test (FET) for the likelihood of site-wise support of "Non-Concerted" and "Concerted" topologies.**

| | # sites | Non-Concerted topology | Concerted topology | FET p-value |
|---|---|---|---|---|
| Nonsynonymous | 28 | 17 | 11 | 3.2e-8 |
| Synonymous | 164 | 9 | 153 | |

**Table S5. List of primers used for intron sequencing**

| Species | Primer |
|---|---|
| *Engystomops pustulosus* | N-terminal<br>Forward: EP_wwBC6_1F<br>GATGTAGAGGGTACGGTTTGAGGCACATGGCGGCAAGAAGAA<br>Reverse: EP_wwBC6_11R<br>GATGTAGAGGGTACGGTTTGAGGCGTGGAGCATCGGTCCAGGA<br>C-terminal<br>Forward: EP_wwBC7_11F<br>GGCTCCATAGGAACTCACGCTACTGATCCTGGACCGATGCTCCA<br>Reverse: EP_wwBC7_19R<br>GGCTCCATAGGAACTCACGCTACTTGACAATGCTGACGAAGAAGGC |
| *Lithodytes lineatus* | Forward: Lep_wwBC3_1F<br>TACATGCTCCTGTTGTTAGGGAGGACATGGCGGCAAGAAGAA<br>Reverse: Lep_wwBC3_21R<br>TACATGCTCCTGTTGTTAGGGAGGAGGCACAGAACCACCATGT |
| *Leptodactylus pentadactylus* | Forward: Lep_wwBC5_1F<br>ACAGCATCAATGTTTGGCTAGTTGACATGGCGGCAAGAAGAA<br>Reverse: Lep_wwBC5_21R<br>ACAGCATCAATGTTTGGCTAGTTGAGGCACAGAACCACCATGT |
| *Leptodactylus latrans* | Forward: Lep_wwBC2_1F<br>AGGTGATCCCAACAAGCGTAAGTAACATGGCGGCAAGAAGAA<br>Reverse: Lep_wwBC2_21R<br>AGGTGATCCCAACAAGCGTAAGTAAGGCACAGAACCACCATGT |
| *Leptodactylus insularum* | Forward: Lep_wwBC1_1F<br>AACGGAGGAGTTAGTTGGATGATCACATGGCGGCAAGAAGAA<br>Reverse: Lep_wwBC1_21R<br>AACGGAGGAGTTAGTTGGATGATCAGGCACAGAACCACCATGT |
| *Leptodactylus colombiensis* | Forward: Lep_wwBC8_23F<br>AGAGGGTACTATGTGCCTCAGCACAAGTATGAGCCCGCAGCCACTTC<br>Reverse: Lep_wwBC8_3044R<br>AGAGGGTACTATGTGCCTCAGCACCCAGGGCTGCGTCTGATGATTAA |

**Table S6. List of engineered ATP1A1 constructs used to test functional effects of amino acid substitutions in *Leptodactylus*.**

| Construct Name | Engineered Substitution(s) | Description |
|---|---|---|
| S | - | Sensitive (S) paralog of *L. latrans* ATP1A1 |
| S+Q111R | Q111R | Q111R on the S paralog background |
| S+N122D | N122D | N122D on the S paralog background |
| S+Q111R+N122D | Q111R + N122D | Q111R and N122D on the S paralog background |
| S+10subs | A112T, E116D, I135V, L180Q, I199L, I279V, S403C, L536M, Q701L, I788M | All substitutions strongly distinguishing R and S paralogs, except Q111R and N122D, on the S paralog background |
| R–Q111R–N122D | R111Q, D122N | Reversions R111Q and D122N on the R paralog background |
| S+12subs | Q111R, A112T, E116D, N122D, I135V, L180Q, I199L, I279V, S403C, L536M, Q701L, I788M | Twelve substitutions strongly distinguishing R and S paralogs on the S paralog background |
| R | - | Resistant (R) paralog of *L. latrans* ATP1A1 |

Note: R and S paralogs of *L. latrans* differ by the 12 substitutions that are the focus of this study and by 9 additional amino-acid substitutions and a two-amino acid insertion-deletion difference. Our experiments revealed that these 10 *latrans*-specific substitutions do not contribute detectably to S vs. R differences in CG resistance of enzyme function (using all 10 as one co-variate, ANOVA p>0.5. Following convention, positions of substitutions are standardized relative to the sheep (*Ovis aries*) sequence NM_001009360 - 5 AA from 5' end.

**Table S7. Summary of the ouabain sensitivity and catalytic properties of $Na^{+}$,$K^{+}$-ATPase for each ATP1A1 construct.** The values represent the mean and standard deviation (SD) ouabain sensitivity ($log_{10}IC_{50}$) of ATPase activity of six biological replicates. ATP1B1 of *Leptodactylus latrans* was co-expressed with ATP1A1.

| ATP1A1 Constructs | Ouabain sensitivity (mol/L) Mean(log10 $IC_{50}$) ± SD | ATPase activity nmol Pi/(mg protein*min) ± SD |
|---|---|---|
| S | -5.63 ± 0.59 | 16.30 ± 5.01 |
| S+Q111R | -4.89 ± 0.85 | 10.68 ± 3.71 |
| S+N122D | -5.06 ± 0.66 | 7.16 ± 3.62 |
| S+Q111R+N122D | -3.62 ± 0.28 | 8.29 ± 3.86 |
| S+10subs | -5.82 ± 0.47 | 16.37 ± 3.05 |
| R–Q111R– N122D | -5.60 ± 0.33 | 17.41 ± 2.87 |
| S+12subs | -3.23 ± 0.75 | 12.17 ± 2.43 |
| R | -3.25 ± 0.77 | 14.09 ± 2.77 |

**Table S8. Statistical analysis of ouabain sensitivity and ATPase activity.** Significant p values are highlighted in bold.

| **(Explanatory Variables) ANOVA** | Ouabain sensitivity log10($IC_{50}$) | | | | ATPase activity nmol Pi/(mg protein*min) | | | |
|---|---|---|---|---|---|---|---|---|
| | df | MS | F | p value | df | MS | F | p value |
| Q111R | 1, 42 | 42.8 | 107.8 | **2.7e-13** | 1, 42 | 83.0 | 6.9 | **0.015** |
| N122D | 1, 42 | 11.8 | 27.72 | **2.3e-6** | 1, 42 | 101.4 | 7.98 | **7.2e-3** |
| 10subs | 1, 42 | 0.59 | 1.6 | 0.22 | 1, 42 | 228.1 | 17.96 | **1.2e-4** |
| R-S background | 1, 42 | 0.04 | 1.9 | 0.74 | 1, 42 | 11.5 | 0.34 | 0.34 |
| Q111R:N122D | - | - | - | - | 1, 42 | 7.6 | 5.64 | **0.022** |

| **(Explanatory Variables) Linear regression** | Ouabain sensitivity log10($IC_{50}$) | | | | ATPase activity nmol Pi/(mg protein*min) | | | |
|---|---|---|---|---|---|---|---|---|
| | Est | SE | t | p value | Est | SE | t | p value |
| Intercept | -6.03 | 0.17 | -36.3 | **<2e-16** | 14.3 | 1.19 | 12.1 | **3e-15** |
| Q111R | 1.32 | 0.21 | 6.27 | **2.7e-13** | -4.39 | 1.88 | -2.34 | **0.024** |
| N122D | 1.14 | 0.21 | 5.45 | **2.3e-6** | -6.81 | 1.88 | -3.63 | **7.7e-4** |
| 10subs | 0.26 | 0.22 | 1.19 | 0.24 | 1.94 | 1.46 | 17.96 | 0.18 |
| R-S background | -0.08 | 0.26 | -0.34 | 0.74 | 1.39 | 1.46 | 0.34 | 0.35 |
| Q111R:N122D | - | - | - | - | 6.90 | 2.91 | 5.64 | **0.022** |

Note: "R-S background" in the ANOVA refers to 9 additional amino acid substitutions and a two amino acid insertion-deletion difference that distinguishes the R and S constructs (derived from *Leptodactylus latrans*).

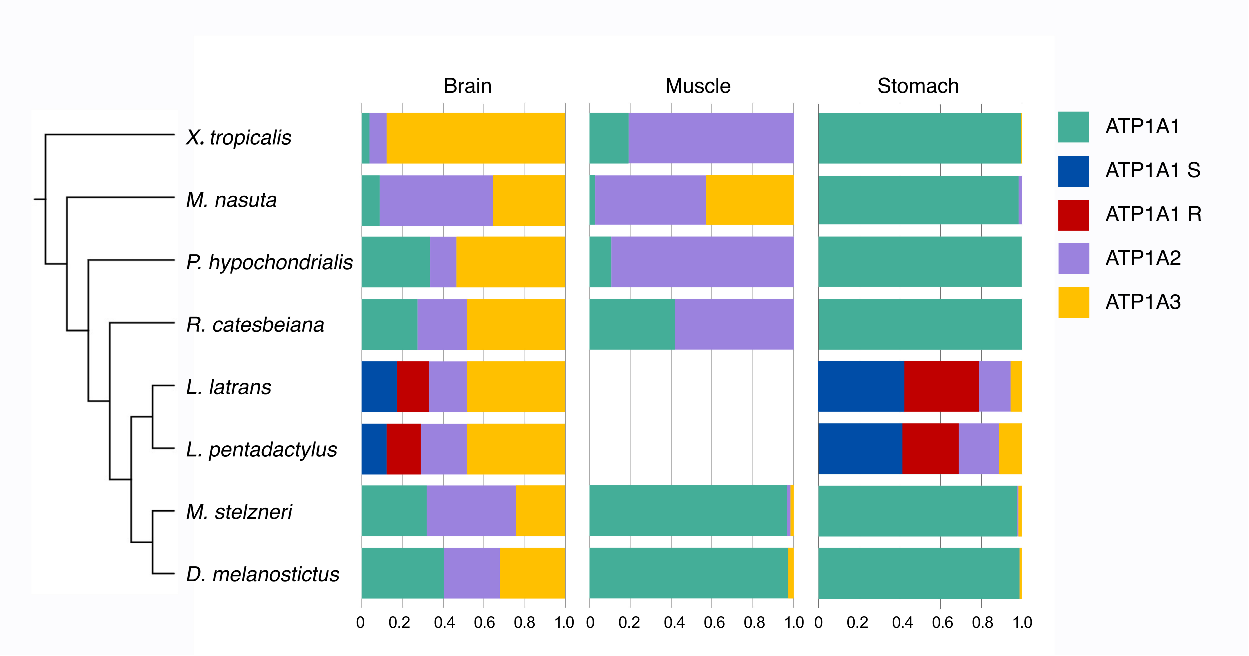


**Fig S1. Proportion of ATP1A1, ATP1A2, and ATP1A3 paralogs in brain, muscle, and stomach of seven anuran species.** RNA-seq reads for eight species were mapped to species-specific copies of ATP1A1, ATP1A2, and ATP1A3 using bwa (see Methods). Uniquely mapped reads were counted for each paralog and estimated as a proportion of the sum of the reads for all three ATP1A paralogs. *X. tropicalis*: *Xenopus tropicalis*; *M. nasuta*: *Megophrys nasuta*; *P. hypochondrialis*: *Phyllomedusa hypochondrialis*; *R. catesbeiana*: *Rana catesbeiana*; *L. latrans*: *Leptodactylus latrans*; *L. pentadactylus*: *Leptodactylus pentadactylus*; *M. stelzneri*: *Melanophryniscus stelzneri*; *D. melanostictus*: *Duttaphrynus melanostictus*.

| Family | Species | ATP1A1 | | | | | | | | | | ATP1A2 | | | | | | | | | | ATP1A3 | | | | | | | | | |
|---|---|---|---|---|---|---|---|---|---|---|---|---|---|---|---|---|---|---|---|---|---|---|---|---|---|---|---|---|---|---|---|
| | | 108 | 111 | 112 | 114 | 115 | 116 | 117 | 119 | 120 | 122 | 108 | 111 | 112 | 114 | 115 | 116 | 117 | 119 | 120 | 122 | 108 | 111 | 112 | 114 | 115 | 116 | 117 | 119 | 120 | 122 |
| | | Y | Q | A | T | E | E | E | Q | N | N | Y | Q | A | M | E | D | E | Q | N | N | Y | Q | A | T | E | D | D | S | G | N |
| Human | *Homo sapiens* | · | · | · | · | · | · | · | · | · | · | · | · | · | · | · | · | · | S | · | · | · | · | · | · | · | · | · | · | · | · |
| Brown rat | *Rattus norvegicus* | · | R | S | · | · | · | · | P | · | D | · | L | · | · | · | · | · | S | · | · | · | · | · | · | · | · | · | · | · | · |
| Bombinatoridae | *Bombina maxima* | · | · | · | · | · | D | · | · | · | · | | | | | | | | | | | | | | | | | | | | |
| Pipidae | *Xenopus tropicalis* | · | T | · | · | · | · | · | T | · | · | · | · | I | · | · | · | · | I | · | · | · | L | · | M | · | E | E | · | · | · |
| Pelobatidae | *Pelobates fuscus* | · | · | · | · | · | · | · | · | · | · | · | · | I | · | · | · | · | I | · | · | · | · | · | · | · | · | · | A | · | · |
| Megophryidae | *Megophrys nasuta* | · | · | · | · | · | · | · | N | · | · | · | · | · | · | · | · | · | · | · | · | · | · | · | · | · | · | · | A | · | · |
| Megophryidae | *Oreolalax rhodostigmatus* | · | · | · | · | · | · | · | · | · | · | · | · | I | · | · | · | · | I | · | · | · | · | · | | | | | | | |
| Megophryidae | *Leptobrachium boringii* | · | · | · | · | · | · | · | · | · | · | | | | | | | | | | | · | · | · | · | · | · | · | A | · | · |
| Microhylidae | *Kaloula pulchra* | · | · | · | · | · | · | D | · | · | · | · | · | I | · | · | · | · | I | · | · | | | | | | | | | | |
| Mantellidae | *Mantella betsileo* | · | · | · | · | · | · | · | · | · | · | | | | | | | | | | | · | · | I | M | · | · | E | I | N | · |
| Dicroglossidae | *Quasipaa boulengeri* | · | · | · | · | · | D | · | · | · | · | | | | | | | | | | | | | | | | | | | | |
| Dicroglossidae | *Fejervarya cancrivora* | · | · | · | · | · | · | D | · | · | · | · | · | I | · | · | · | · | I | · | · | | | | | | | | | | |
| Ranidae | *Pelophylax lessonae* | · | · | · | · | · | · | · | · | · | · | · | · | I | · | · | · | · | I | · | · | · | · | · | · | · | · | · | A | N | · |
| Ranidae | *Odorrana tormota* | · | · | · | · | · | · | · | · | · | · | · | · | I | · | · | · | · | I | · | · | · | · | · | M | · | · | · | A | N | · |
| Ranidae | *Rana sphenocephala* | · | · | I | M | · | D | · | I | · | · | · | · | I | · | · | · | · | I | · | · | · | L | · | · | · | · | · | · | N | · |
| Ranidae | *Rana catesbeiana* | · | · | · | · | · | · | · | · | · | · | · | · | I | · | · | · | · | I | · | · | · | L | · | · | · | · | · | · | N | · |
| Myobatrachidae | *Limnodynastes peronii* | · | · | · | · | · | · | · | · | · | · | · | · | I | · | · | · | · | I | · | · | · | · | · | · | · | · | · | A | N | · |
| Hylidae | *Cyclorana alboguttata* | · | · | · | · | · | · | · | · | · | · | · | · | I | · | · | · | · | I | · | · | | | | | | | | | | |
| Hylidae | *Phyllomedusa hypochondrialis* | · | · | · | · | · | · | · | · | · | · | · | · | I | · | · | · | · | I | · | · | · | · | · | · | · | · | · | A | · | · |
| Craugastoridae | *Craugastor fitzingeri* | · | · | · | · | · | · | · | · | · | · | · | · | I | · | · | · | · | I | · | · | | | | | | | | | | |
| Strabomantidae | *Oreobates cruralis* | · | · | · | · | · | · | · | · | · | · | · | · | I | · | · | · | · | I | · | · | | | | | | | | | | |
| Leptodactylidae | *Engystomops pustulosus* | · | · | · | · | · | · | · | · | · | · | · | · | I | · | · | · | · | I | · | · | · | · | · | · | · | · | · | A | · | · |
| Leptodactylidae | *Lithodytes lineatus* | · | · | · | · | · | · | D | · | · | · | · | · | I | · | · | · | · | I | · | · | | | | | | | | | | |
| Leptodactylidae | *Leptodactylus latrans* S | · | · | · | · | · | · | · | · | · | · | · | · | I | · | · | · | · | I | · | · | · | · | · | · | · | · | · | A | · | · |
| | *Leptodactylus latrans* R | · | R | T | · | · | D | · | · | · | D | | | | | | | | | | | | | | | | | | | | |
| Dendrobatidae | *Dendrobates auratus* | · | · | · | · | · | · | · | · | · | · | · | · | I | · | · | · | · | I | · | · | · | · | · | · | · | · | · | T | · | · |
| Bufonidae | *Melanophryniscus stelzneri* | H | L | V | · | · | D | · | N | · | · | · | · | I | · | · | · | · | M | · | · | · | · | · | · | · | · | N | A | N | · |
| Bufonidae | *Atelopus zeteki* | · | R | K | S | D | L | · | D | · | · | | | | | | | | | | | | | | | | | | | | |
| Bufonidae | *Duttaphrynus melanostictus* | · | R | K | S | D | L | · | D | · | · | · | T | V | I | · | · | D | T | · | · | · | L | · | · | · | · | E | · | R | · |
| Bufonidae | *Bufotes viridis* | · | R | K | S | D | L | · | D | · | · | · | T | V | I | · | · | D | T | · | · | · | R | K | S | D | L | E | D | N | · |
| Bufonidae | *Rhinella marina* | · | R | K | S | D | L | · | D | · | · | | | | | | | | | | | · | L | · | · | · | · | E | · | R | · |

**Fig S2. Variation among sites implicated in CG-resistance for ATP1A paralogs of various species.** Sequences of ATP1A2 and ATP1A3 were reconstructed using the same method as ATP1A1 described in Materials and Methods. Consensus sequences of anuran species were generated in MEGA 7.0 and used as reference for each paralog. Only sites implicated in CG-resistance are shown. Following convention, positions of substitutions, shown at the top, are aligned relative to the sheep (*Ovis aries*) sequence NM_001009360 subtracting 5 AA from 5'end (e.g., the first position is 108). A dot indicates identity with the reference sequence. ATP1A1S and ATP1A1R of *Leptodactylus latrans* are indicated in blue and red, respectively. Bufonid (toad) species, the prey species that produce CG toxins, are highlighted in purple. Blank: missing data. We failed to identify an ortholog of ATP1A4 in any of the available anuran genome assemblies, including our assembly of *Leptodactylus fuscus*.

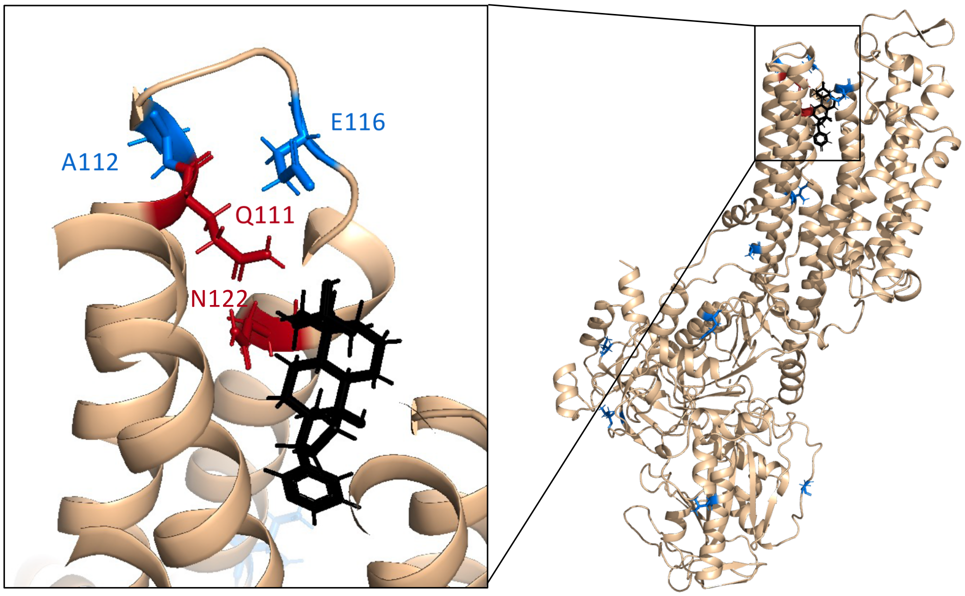


**Fig S3. Positions of 12 R copy-specific amino acid substitutions on the crystal structure of pig ATP1A1 bound to the cardiac glycoside bufalin.** Red residues correspond to key CG resistance-conferring sites 111 and 122; blue residues correspond to 10 additional residues strongly associated with variants at sites 111 and 122. (*Sus scrofa*, PDB 4RES.)

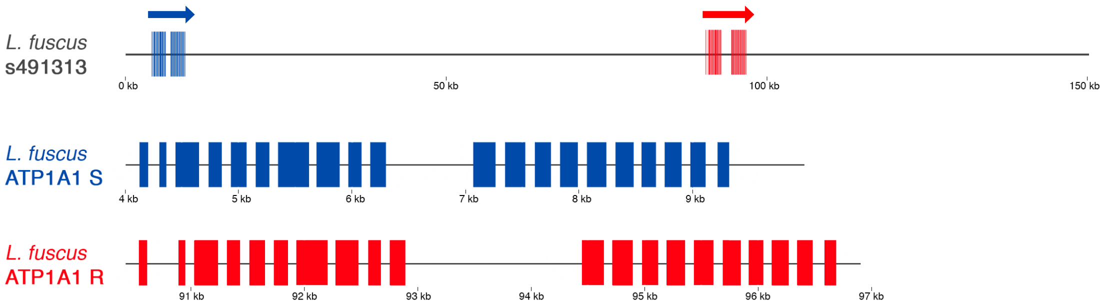


**Fig S4. Annotation of ATP1A1S and ATP1A1R paralogs in the *Leptodactylus fuscus de novo* genome assembly (Table S3)**. ATP1A1S (blue) and ATP1A1R (red) occur in tandem on scaffold s491313 (Genbank Acc#) ~80 kilobases apart. The boundary between exons and introns were determined by blast and manual correction (*i.e*., ensuring that each intron started with GT and ended with AG). The gene structure figures were plotted with ggbio in R.

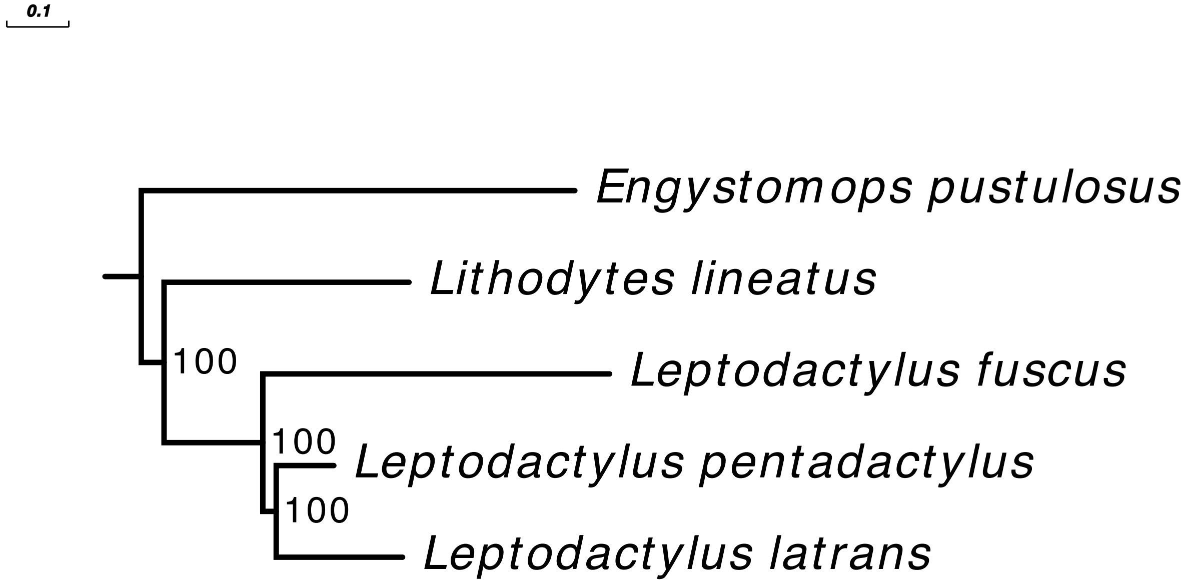


**Figure S5. A phylogenetic tree of *Leptodactylus* and outgroup species.** The tree was constructed using an alignment of 813 orthologous mRNA sequences under the best partition model with iq-tree 2.0 (see Methods). Branch lengths: (*Engystomops pustulosus*:0.699436, (*Lithodytes lineatus*:0.395135, (*Leptodactylus fuscus*:0.559378, (*Leptodactylus pentadactylus*: 0.092965, *Leptodactylus latrans*: 0.203845)100:0.0216082)100:0.159296)100:0.0368124).

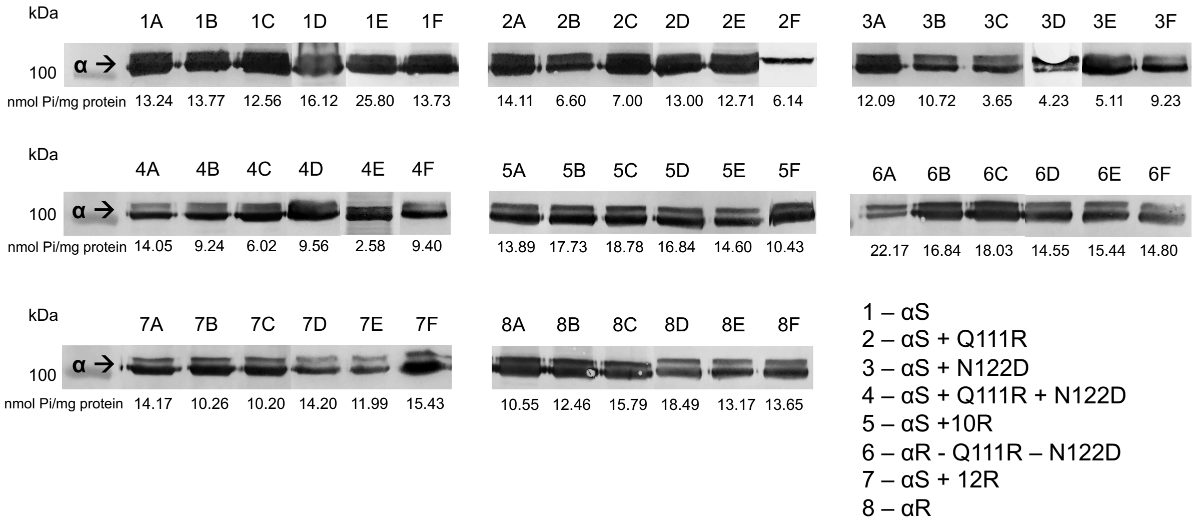


**Fig S6. Western blot analysis of Na$^+$,K$^+$-ATPase with engineered ATP1A1 (α) subunits produced in this study.** The 110 kDa ATP1A1 protein is stained with the α5 monoclonal antibody followed by a horseradish peroxidase conjugated goat antimouse antibody. Samples represent six biological replicates of eight different recombinant Na$^+$,K$^+$-ATPase (Table S5) produced through cell culture. All western blots are aligned to the same protein ladder level. ATPase activity levels (nmol Pi/mg protein) of each biological replicate is indicated under its respective band.

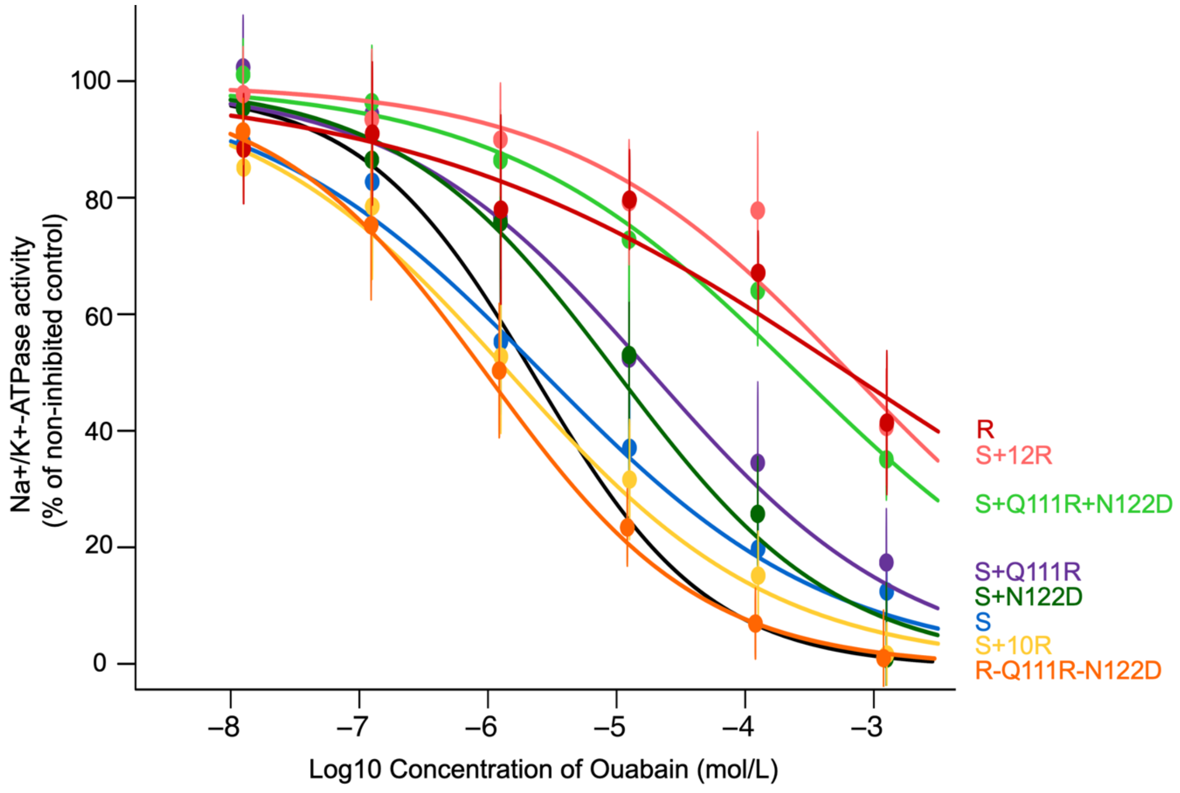


**Fig S7. Cardiac glycoside (ouabain) inhibition curves for six each engineered *Leptodactylus* Na$^+$,K$^+$-ATPase produced in this study.** Points and error bars represent the mean ± SEM (n=6 biological replicates) percentage of protein activity relative to controls measured in the absence of ouabain and excluding the activity of background ATPases. The black inhibition curve was measured from commercially procured porcine cerebral cortex (CAS 9000-83-3, Sigma-Aldrich, Inc.) and represents a standard bench-mark reference for cardiac glycoside sensitivity (Log10 $IC_{50}$= -5.61). The ATP1B1 of *Leptodactylus latrans* was co-expressed with each engineered version of ATP1A1 (Table S5).
